# Circulating pre-osteoclasts are primed for osteoclast fate and synovial tissue homing in psoriatic arthritis

**DOI:** 10.64898/2026.03.21.713366

**Authors:** Joseph Hutton, Atrik Saha, Qian Li, Neda Farahi, Francesca Leek, Stephen M McDonnell, Busola Ade-Ojo, Daniel Gillett, Nicholas Shenker, Deepak Jadon, Anna Laddach, Charlotte Summers, Naomi McGovern

## Abstract

Psoriatic arthritis (PsA) is a complex immune-mediated inflammatory disease with heterogenous clinical features. Osteoclasts have a unique ability to destroy bone, playing key roles in both healthy bone turnover and pathological erosions in arthritis. They are believed to arise from monocytic precursors migrating to inflamed synovial tissue, though the identity of this precursor in humans has remained elusive. Here, we sought to determine whether monocytes home to psoriatic joints and their phenotype. We find that monocytes are recruited to inflamed joints in PsA but not uninvolved or healthy joints, and identify a pre-osteoclast (preOC) cell state derived from classical monocytes. PreOC are transcriptionally distinct from classical monocytes, are enriched for *NFACT1*, the master regulator of osteoclastogenesis, and form multinucleated osteoclasts *in vitro* without exogenous RANKL. They are expanded in the circulation of donors with active treatment naïve PsA, compared to both healthy controls and those with cutaneous psoriasis only. Finally, using a customised spatial transcriptomics workflow, we identified these cells within human synovial tissue, and radiolabelling confirmed their recruitment from the circulation. These findings highlight the role of systemic immune priming in macrophage pathophysiology.

## Introduction

Psoriasis is a common immune-mediated skin condition that affects about 1 in 50 people in the United Kingdom (UK)^1^. Approximately one-third of patients will develop psoriatic arthritis (PsA)^1^, a common immune-mediated inflammatory disease. PsA is a heterogenous condition with diverse manifestations including peripheral inflammatory arthritis (IA), axial inflammation, enthesitis, nail and skin disease, and extra-articular manifestations like gut and eye involvement^2^. Psoriatic synovial disease includes inflammation, neovascularisation, pannus formation, bony erosions, and the phenomenon of marked osteolysis in the arthritis mutilans phenotype.

Osteoclasts are highly specialised macrophages solely responsible for bone resorption during development, healthy bone maintenance, and disease states^3, 4^. They are multinucleated giant cells formed by the fusion of multiple cells of both primitive and definitive haematopoietic origins^4, 5^. Which exact definitive haematopoietic cells contribute to osteoclastogenesis has remained ambiguous, despite the established ability of various myeloid cells, particularly monocytes, to form osteoclasts^6–8^. This uncertainty is underscored by high interdonor variability in osteoclastogenic potential *in vitro*^8^ and considerable technical challenges of studying these cells *in vivo* in humans^9^. The precise markers defining osteoclast-biased populations continue to be debated^9–12^. Whilst promising candidate populations, such as the DCSTAMP^pos^ (dendritic cell specific transmembrane protein) cells^11, 13, 14^, have been identified, these have lacked the in-depth characterisation needed to definitively establish their role as a distinct lineage.

Rheumatoid arthritis (RA) shares many clinical features with PsA, including bony erosion, yet has significant differences in its clinical, genetic and cellular-molecular drivers. The mechanistic disparities are critical in driving their unique manifestations and differential response to disease-modifying therapies.

In IA, there is abnormal leukocyte trafficking, with circulating cells possessing an inherently pro-migratory and adhesive phenotype^15, 16^. Although therapeutically targeting leukocyte trafficking pathways has succeeded in other inflammatory diseases, it has had limited success in PsA and other IA^17, 18^. Understanding the specific cellular interactions driving aberrant recruitment in PsA remains a critical, unresolved challenge^17, 18^.

Synovial tissue macrophages are a prominent cell type in PsA synovium, and PsA features similar levels of CD14^pos^ macrophages to RA, believed, but not confirmed to be, reflective of recruitment of monocytes to joint tissue^2, 19^. As such, and given monocyte importance to osteoclastogenesis, we sought to establish whether monocytes are specifically recruited to psoriatic joints. Alongside this, we sought to characterise the tissue homing phenotype of mononuclear phagocytes in a systematic unbiased manner, to best identify potentially relevant markers for synovial tissue homing in active treatment naïve PsA. Here, we determine using radiolabelled techniques that monocyte recruitment to inflamed psoriatic joints is increased in comparison to uninflamed joints or those of healthy controls. We identify a range of tissue homing markers differentially expressed between health and disease, and a new cell state, termed pre-osteoclasts (preOC). PreOC are transcriptionally distinct from classical monocytes, closely resemble *in vitro* generated monocyte-osteoclast intermediaries, and are enriched for osteoclastogenesis and osteoclast-function genes, including *NFATC1*. PreOC are expanded in the peripheral blood of patients with active treatment naïve PsA compared to both healthy and cutaneous psoriasis controls. We establish *in vitro* culture assays for sort purified preOC, and find that, in comparison to classical monocytes, they can form osteoclasts in the absence of exogenous RANKL, the key osteoclast inducing factor. Both circulating and synovial preOC express a synovial tissue homing signal and we identify an expansion of preOC within psoriatic synovium compared to healthy and other inflammatory arthritis synovium. *In silico* analyses reveal a preOC to osteoclast trajectory within synovial tissue. Finally, we use imaging mass cytometry to trace and phenotype radiolabelled circulating cells into tissue, and demonstrate enrichment of this signal in osteoclast-lineage cells.

This study demonstrates that the osteoclast fate can be pre-determined before precursors reach the joint. Additionally, it provides insight into how abnormal preOC homing fuels structural damage in PsA, highlighting the role of systemic monocyte priming in disease pathophysiology.

## Results

### Monocytes demonstrate increased recruitment to inflamed joints in patients with active psoriatic arthritis

Tissue macrophages are expanded in abundance in PsA synovium^2, 19, 20^. We sought to establish if increased monocytic recruitment contributes to this expansion. Monocyte-specific radiolabelling has been used to show uptake of monocytes to joint tissue, though not previously in PsA patients^21–23^.

We utilised technetium^99m–^HMPAO radiolabelling of autologous monocytes (Tc^99m–^monocytes) and planar imaging to characterise their kinetics and migration to key sites of interest (hands and wrists, knees, feet and ankles) in healthy volunteers and participants with active PsA (**Figure 1A**). Baseline demographics of participants are included in **Supplementary Table 1**. Briefly, all participants with PsA met CASPAR-criteria, and had active erosive disease (**Supplementary Table 2**). Details of monocyte radiolabelling parameters are provided in **Supplementary Table 3**. In healthy volunteers, Tc^99m–^monocytes are not specifically detected in joints over the sequential planar imaging (**Figure 1B**). In contrast, Tc^99m–^monocytes uptake is observed in the clinically inflamed joints of participants with active PsA, but not in their non-inflamed joints (**Figure 1C**). Sequential imaging demonstrated an increase in the uptake of Tc^99m–^monocytes within the wrist joint regions of interest (ROIs) in all participants with active PsA. Conversely, signal intensity in healthy controls progressively declined over the same period, suggesting an absence of active monocyte recruitment (**Figure 1D**). Monocyte uptake to inflamed joints correlates with evidence of active synovitis with effusions, hyperaemia and synovial proliferation on contemporaneous musculoskeletal ultrasound (**Figure 1E, Supplementary Figure 1**).

**Figure 1.**
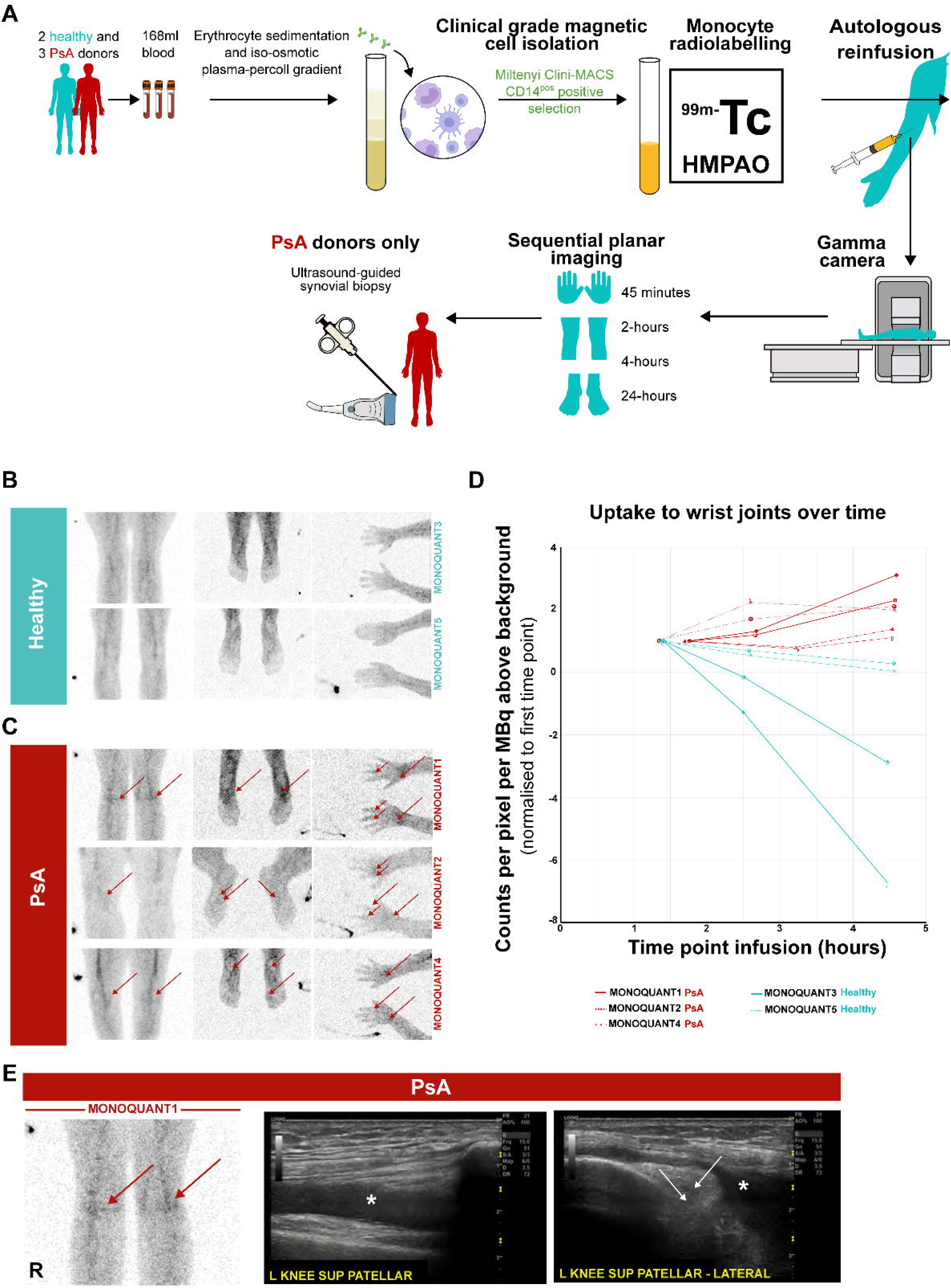
Monocytes demonstrate increased recruitment to inflamed joints in patients with active psoriatic arthritis. **A)** Schematic of workflow for radiolabelling imaging studies. 2 healthy volunteers and 3 patients with active PsA were recruited. 168ml of blood was taken and clinical grade PBMCs isolated using erythrocyte sedimentation and iso-osmotic plasma-percoll gradients. Monocytes were selected using magnetic positive selection using CliniMACS CD14^pos^ beads. Purified monocytes were labelled with a target of 1000MBq of Tc^99m–^HMPAO, and autologously reinfused. Initial dynamic planar imaging was performed over the lungs and upper abdomen to explore their reinfusion kinetics. Following this, sequential planar images of participants knees, feet/ankles, and hands/wrists performed at 45-minutes, 2-, 4-, and 24-hours. Donors with PsA underwent an ultrasound-guided synovial biopsy after their 24-hour imaging timepoint. **B, C)** Images of knees, feet/ankles, and hands/wrists from healthy donors (B) and PsA donors (C). Images are composites, representing the geometric mean of counts measured from both anterior and posterior scanners over the 20 minutes of image capture. An exception is that of the hands/wrists which are from posterior scanner only. Right side of image indicated with reference source. Images are normalised to injected activity. Red arrows indicate accumulation of activity inflamed joints. For MONOQUANT1 donor, uptake seen in both knees, ankles, wrists, and right (R) metacarpophalangeal (MCP) 2, and 3, and left (L) MCP 3 joints. For MONOQUANT2 donor, uptake seen in R knee, R metatarsophalangeal (MTP) 1, L MTP 1-2, R wrist, R MCP 1-2, L MCP 1-2 joints. For MONOQUANT4 donor, uptake seen in both knees, ankles, R MTP 1, both wrists, and R MCP1 joints. **D)** Distribution of radioactivity to bilateral wrist joints over time for all donors in counts per pixel per MBq above background radiation, normalised to first time point of imaging. **E)** Correlation of radiolabelling findings with musculoskeletal ultrasound. Top panels indicate inflamed joints of note from PsA donor (MONOQUANT1) who underwent musculoskeletal ultrasound assessment prior to synovial biopsy. To right are relevant ultrasound images of regions. Effusions indicated with *, white arrows indicate errors of synovial hyperplasia. Red arrows indicate accumulation of activity inflamed joints.

### Psoriatic classical monocytes display aberrant expression of tissue homing markers and osteoclastogenesis

Having established that monocytes migrate into inflamed psoriatic joints, we next sought to identify differences in their properties that might govern the tissue homing of mononuclear phagocytes, in an unbiased manner. For this, we sort-purified classical monocytes (cMos), intermediate monocytes (iMos), non-classical monocytes (ncMos), DC1, DC2, DC3, pDC and preDCs from healthy and active treatment naïve PsA blood (**Figure 2A**) using optimised gating strategies (**Supplementary Figure 2A and 2B**). We performed high sequencing depth bulk RNA-seq to a depth of 90 million paired reads on purified populations, to better characterise subtle differences and capture chemokine / adhesion receptors. CD14^pos^ CD163^pos^ DC3, CD5^pos^ DC2s and CD14^neg^ CD163^neg^ DCs were isolated and sequenced as separate populations. However, we subsequently combined both CD5^pos^ DC2s and CD14^neg^ CD163^neg^ DCs into a single DC2 group as their PCA embeddings and marker expression profiles demonstrate these cells have similar transcriptomes, and likely represent a single population (**Supplementary Figure 2C**), in agreement with the findings of others^24–27^. Cell subsets cluster as expected in PCA space (**Supplementary Figure 2C**). Final validation of our sort strategies was performed by assessing for enrichment of subset-specific gene signatures derived from publicly available mononuclear phagocyte scRNA-seq datasets^27, 28^ (**Supplementary Figure 2D**), confirming the identity of our sorted cellular populations.

**Figure 2.**
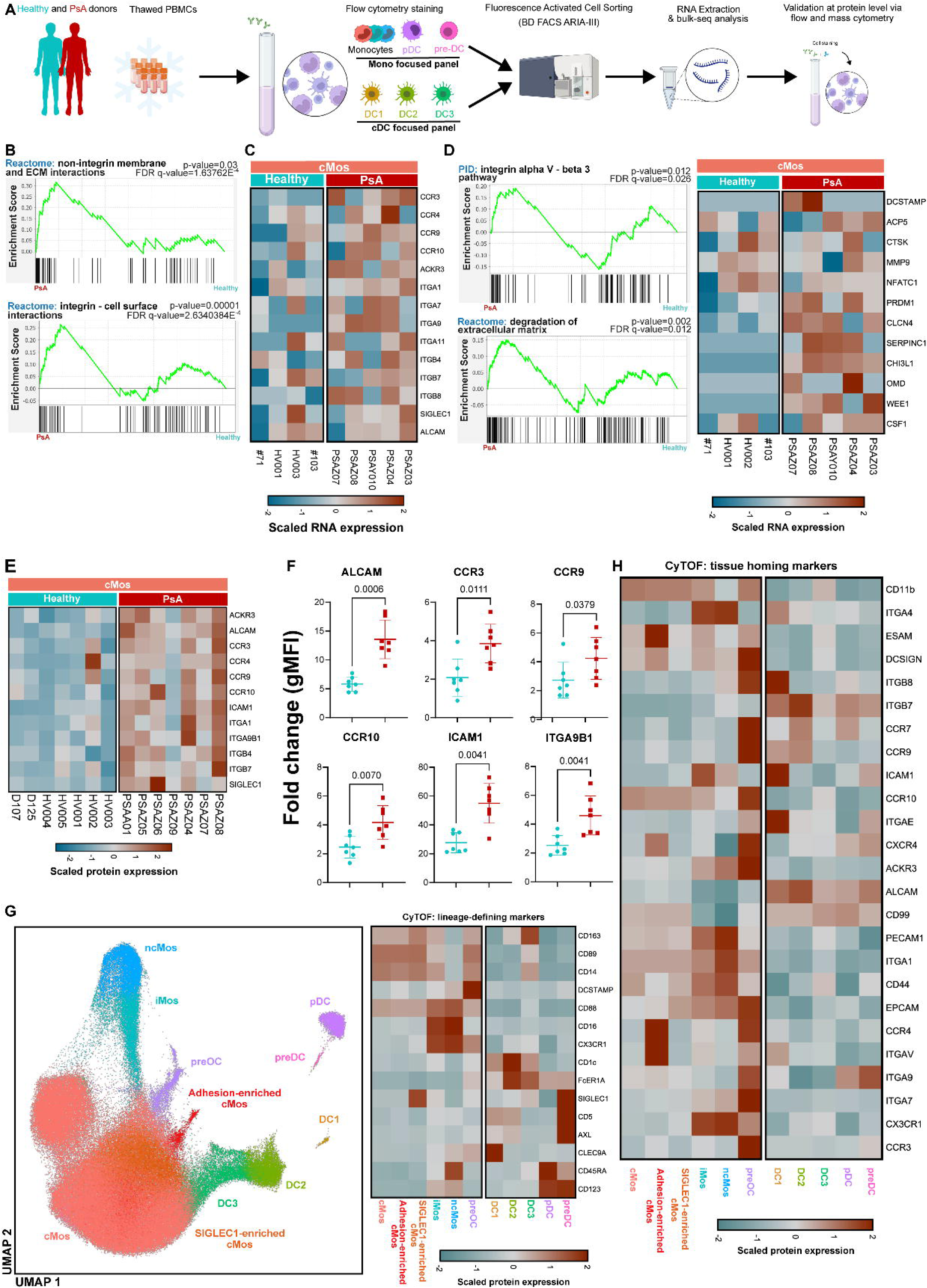
Psoriatic classical monocytes display aberrant expression of tissue homing markers and osteoclastogenesis. **A)** Schematic showing workflow. Frozen PBMCs are thawed, stained with a monocyte- or cDC-focused panel and sorted using the ARIA-III. Cells are sorted directly into RLT lysis buffer and the RNA was immediately extracted. Created with BioRender.com and Inkscape. **B)** Gene-set enrichment analysis plots of key pathways enriched in PsA cMos (upper plot, integrin: cell surface, and lower plot showing non-integrin: extracellular matrix) compared to healthy counterparts. **C)** Heatmap showing scaled gene expression of differentially expressed markers for trafficking and adhesion in cMos in health vs. PsA. **D)** Left panels; gene set enrichment analysis plot demonstrating enrichment of cMos for integrin alpha V – beta 3 (bone binding receptor) and degradation of extracellular matrix gene signature in PsA. Right panel; heatmap showing scaled gene expression of gene markers related to osteoclastogenesis and/or osteoclast function in PsA cMos. **E)** Heatmap showing protein level expression of select markers (scaled expression) via flow cytometry (fold-change gMFI). **F)** Dotplots showing the fold-change in gMFI for select markers compared to FMO for health (cyan) and PsA (red) donors. Mean value with SD are indicated with coloured lines, Mann-Whitney-U test. **G)** Left panel; final UMAP plot of 200,000 blood MPs cells from the 48-parameter mass cytometry panel, identifying key lineages. Populations identified include cMos, iMos, ncMos, DC1, DC2, DC3, preDC, and pDCs. Additional cell states including SIGLEC1-enriched cMos, adhesion-enriched cMos, and putative preOC are also identified. Right panel; heatmap showing scaled expression of mononuclear phagocyte lineage defining markers for the clusters identified. **H)** Heatmap showing scaled expression of tissue homing markers for the clusters identified in G.

To determine how the gene expression profile of circulating mononuclear phagocytes differs between health and PsA, we analysed the differentially expressed genes, focusing on classical monocytes given the demonstrated monocytic uptake to joint tissue. Differences were also observed in DC populations, including enrichment for PsA-relevant cytokines (*IL12A, IL23A, TNF,* and *VEGFA*) in DC2 (**Supplementary Figure 2G**), further supporting the hypothesis that mononuclear phagocytes may be primed even before encountering the joint microenvironment. However, a full exploration of DC was beyond the scope of this study. Through both unsupervised gene set enrichment analysis, and guided exploration of tissue-homing markers, we determine that PsA cMos are enriched for both integrin binding and non-integrin / extracellular matrix binding (**Figure 2B**). Supervised analysis demonstrates enrichment of classical monocytes in PsA for a range of tissue homing markers including *CCR3, CCR4, CCR9, CCR10*, and multiple integrins (**Figure 2C**). Enrichment for both the bone binding receptor ITGAV-ITGB3 and extracellular matrix degradation gene signatures are also observed in cMos in PsA compared to health (**Figure 2D**). A number of genes related to osteoclastogenesis and osteoclast function including *DCSTAMP, ACP5, PRDM1*, and *CLCN4* are enriched in PsA cMos, including strong enrichment of DCSTAMP in 2 donors **(Figure 2D, right heatmap)**.

Differential protein expression of homing markers is confirmed using flow cytometry (gating strategy in **Supplementary Figure 2E**). We observe similar enrichment of these tissue homing markers in psoriatic classical monocytes **(Figure 2E**), with significant enrichment for multiple markers in comparison to healthy classical monocytes (**Figure 2F**).

We sought to further validate these findings by performing proteomic analyses by suspension mass cytometry in health and active treatment naïve PsA peripheral blood mononuclear cells (PBMCs). Contaminating lymphocytes and debris were removed (**Supplementary Figure 2H-I**), samples were down sampled and integrated after batch correction. Uniform manifold approximation and projection (UMAP) visualisation of populations readily identifies known mononuclear phagocyte (MPs) subsets by their well-described canonical markers **(Figure 2G**). It also delineates two additional populations, we tentatively termed pre-osteoclasts (preOCs) and adhesion marker -enriched classical monocytes (adhesion-enriched cMos) (**Figure 2G**). These identities were assigned on the basis of high expression of DCSTAMP (a molecule that is important for cell: cell fusion in osteoclastogenesis^14, 29–32^) **(Figure 2G, right heatmap**), and enrichment of tissue-homing markers on both populations (**Figure 2H**).

### Pre-osteoclasts are enriched in the blood of donors with active treatment naïve PsA compared to healthy and cutaneous psoriasis only controls

UMAP visualisation of the mass cytometry data from healthy controls and participants with active treatment naïve PsA reveals the presence of putative preOCs, missed by standard gating strategies (**Figure 2G**). To further investigate this population, we used HyperFinder to derive a gating strategy to isolate preOC from classical monocytes (**Figure 3A-B**). We applied this gating strategy to healthy controls, active treatment naïve PsA, and cutaneous psoriasis (PsC, psoriasis for at least 10 years duration, no evidence of PsA on both clinical and ultrasonographic assessment by a rheumatologist, naïve to systemic disease modifying therapies) (**Figure 3C-D**). Enumeration of preOCs demonstrates significant enrichment in active treatment naïve PsA compared to both healthy and PsC (**Figure 3E**). One PsC outlier donor illustrated in red later developed PsA (**Figure 3E**). These findings demonstrate that active treatment naïve PsA blood is enriched for circulating preOCs, in comparison to healthy controls and PsC.

**Figure 3.**
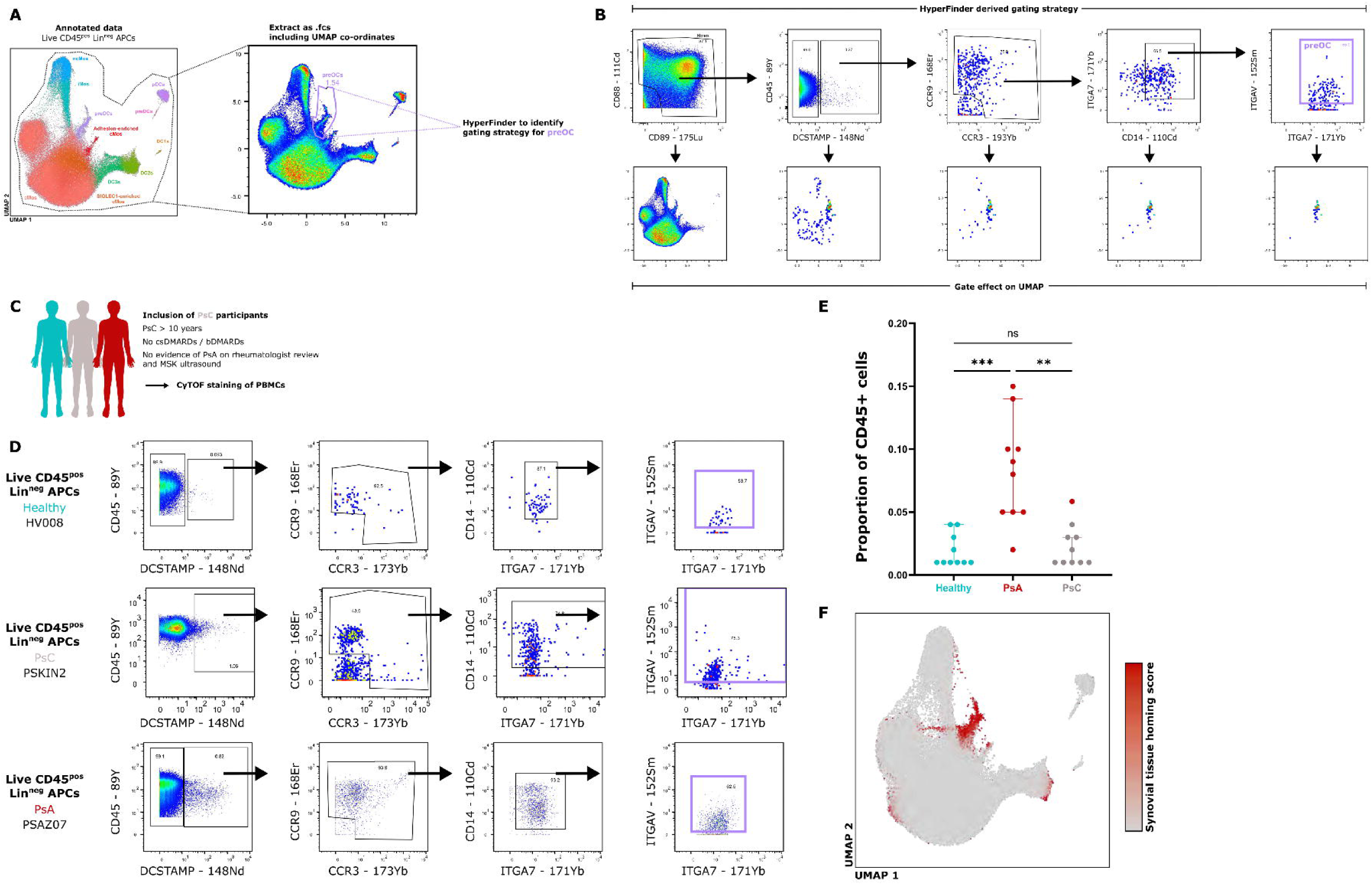
Pre-osteoclasts are enriched in the blood of donors with active treatment naïve PsA compared to healthy and cutaneous psoriasis only controls, and are primed for synovial tissue homing. **A)** Schematic to develop gating strategy for preOC cells. **B)** Output of HyperFinder-derived gating strategy. Top panels show the suggested gating strategy with bottom panels showing the effect of the above gate on the UMAP visualisation. Monocytes are gated on using CD88 and CD89. DCSTAMP^pos^ cells are next selected, already showing strong enrichment for the preOC tail. CCR3^pos^ CCR9^pos^ cells are then gated on, before CD14^pos^ ITGA7^pos^ ITGAV^pos^ cells are selected. **C)** Inclusion and workflow of additional control – cutaneous psoriasis without arthritis (PsC) donors. **D)** Representative HyperFinder-derived gating strategies of healthy (top panels), PsC (middle panels), and PsA (lower panels) of suspension mass cytometry acquired blood PBMCs. **E)** Quantification of preOC population shown as proportion of total CD45^pos^ cells per donor. PsC cohort includes one donor highlighted in red who developed active PsA with inflammatory back pain and dactylitis. P-values were calculated utilising Mann-Witney U, **p-value<0.01, ***p-value<0.001. **F)** UMAP visualisation of blood MPs cells with heatmap overlay of enrichment for “synovial tissue homing” scores from Moon *et al*, 2023.

### PreOCs are primed for a synovial tissue homing signal

Synovial tissue homing markers in RA have been previously identified by others^16, 18, 33–35^. To elucidate if preOC are likely to be recruited to joint tissue, we next sought to assess if the tissue homing markers they express align with these. Several of the tissue homing markers included in our CyTOF panel (CCR3, CCR4, CCR7, CCR10, CXCR4, CX3CR1) overlap with markers important for synovial tissue homing (**Supplementary Figure 3A, Supplementary Table 4**). We find enrichment of this signal in the preOC population (**Figure 3F**). Additionally, PsA preOC express significantly higher levels of multiple tissue homing markers than their counterparts from healthy controls, including CD11b, PECAM1, ICAM1 and CD99, though this does not extend to those overlapping with Moon *et al* (**Supplementary Figure 3B**).

### Pre-osteoclasts are transcriptionally distinct from classical monocytes and resemble *in vitro* generated monocyte – osteoclast intermediaries

To establish if the population identified as preOC represent circulating monocytes that are primed for osteoclast fate (**Supplementary Figure 3C**), we sought to establish their transcriptomic signature and to capture monocyte-preOC intermediaries (**Supplementary Figure 3C**). As circulating preOC are rare, we utilised HyperFinder to further refine the pre-OC gating strategy that would allow us to also sort any transitory cell states between these classical monocytes and preOC (**Supplementary Figure 3D**). Cells were FACS purified accordingly (**Supplementary Figure 3E**). We sequenced 2 healthy and 4 active treatment naïve PsA donors across 3x 384-well plates using VASA-seq scRNA-seq (**Supplementary Figure 3F**). Expression of the key monocytic marker CD14-PE-Dazzle594 was confirmed using CellView Imaging on the BD FACSDiscover S8 at the time of FACS purification (**Supplementary Figure 3G**) as an additional purity check. After pre-processing, quality control (**Supplementary Figure 4A**) and the removal of 67 contaminating B cells (*CD79A, MS4A1*) from the dataset, 658 cells were annotated into 3 clusters based on their gene expression profiles **(Figure 4A**). One population of cMos and 2 populations of preOC (preOC-1, preOC-2) are identified **(Figure 4A**). These populations are present in both health and PsA (**Figure 4B**) and all donors contained cells from each population (**Figure 4C**). There is progressive enrichment of the suspension mass cytometry-identified “preOC” signature across preOC-1 and preOC-2 (**Figure 4D**). This includes progressive enrichment of the master regulator of osteoclastogenesis *NFATC1* in preOC-1 and preOC-2 (**Figure 4E**). No such progressive enrichment is seen for the pan-monocyte marker *C5AR1* (CD88) (**Figure 4F**). Classical monocytes show low levels of enrichment for *NFATC1*, although this is confined to psoriatic cMos only (**Figure 4G**). Analysis of differentially expressed genes (DEGs) between the clusters reveals heterogeneity in the cMos and preOC compartment, with progressive downregulation of monocytic markers (*CD14, S100A8, S100A9*) and upregulation of genes involved in osteoclastogenesis (*NFATC1, NOTCH2, CEBPA, TRAF1, TRAF6, FOS, JUN, SRC, CITED2, FOXM1*) (**Figure 4H**). PreOC also show progressive enrichment for transcripts encoding important aspects of osteoclast biology including proton pumps (*ATPV0A1*, *ATP6V0D1*), proteases (*TCIRG1, CST1, CTSA, ANPEP, SPG7, CA2*), and the bone binding receptor (*ITGAV, ITGB3*) (**Figure 4H**). Finally, preOC-1 and −2 also show enrichment for pro-inflammatory cytokines and chemokines including *CCL3, CCL4, IL1B, TNF, and VEGFA* (**Figure 4H**).

**Figure 4.**
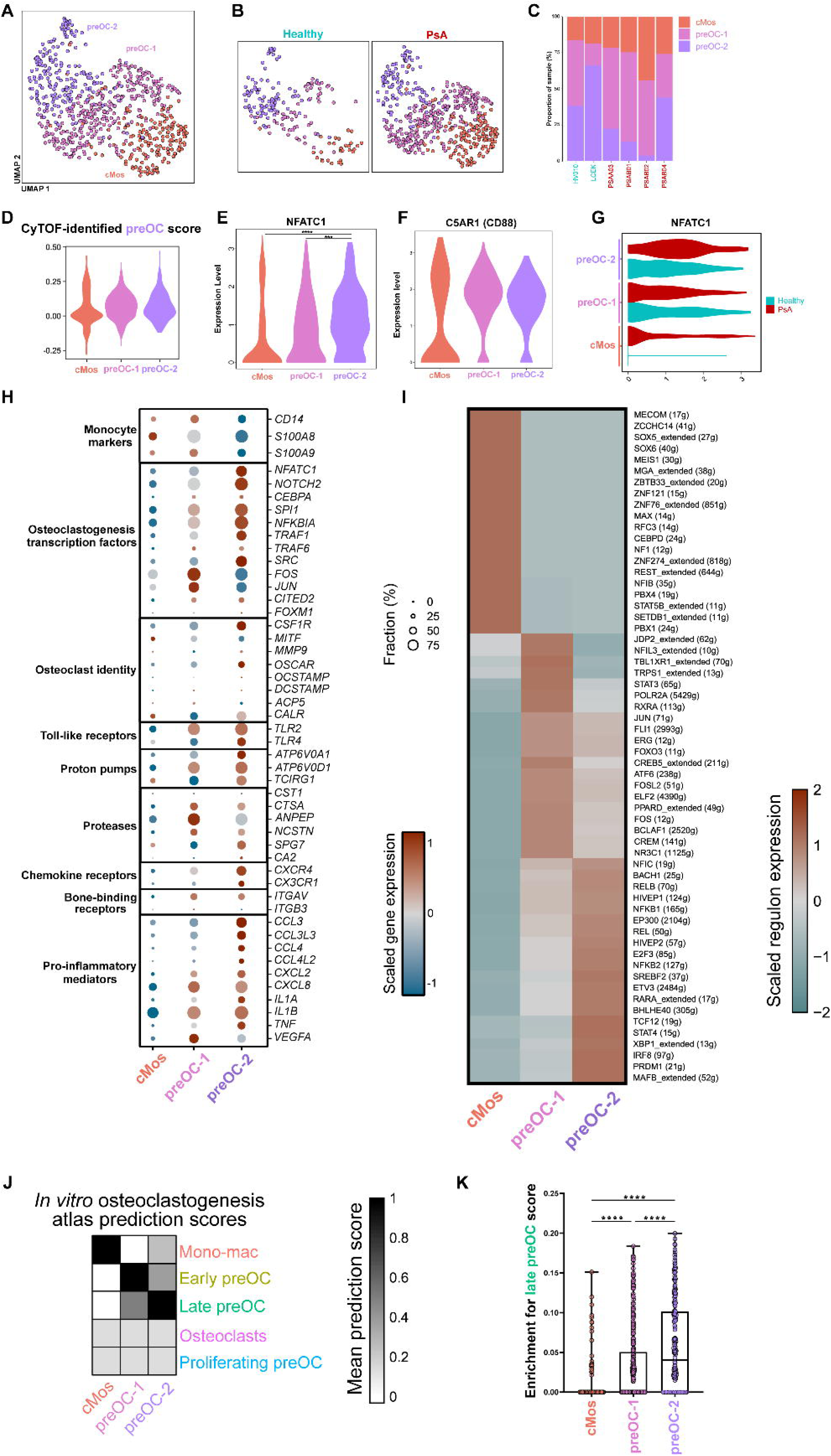
Pre-osteoclasts are transcriptionally distinct from classical monocytes and resemble *in vitro* generated monocyte – osteoclast intermediaries. **A)** UMAP visualisation of 658 cells from 4 active treatment naïve PsA, and 2 healthy donors. Cells are coloured by their final annotation (red-pink cMos, light pink preOC-1, purple preOC-2). **B)** UMAP visualisation showing object split by disease state (health vs. PsA). **C)** Stacked bar chart indicating the proportion of cells from each cluster derived from each donor. **D)** Violin plot showing enrichment for suspension mass cytometry identified markers on plate-based populations via AddModuleScore of CyTOF-derived preOC scores. **E)** Violin plot displaying log-normalised *NFATC1* expression between cMos, preOC-1, and preOC-2 clusters. P-values calculated by 2 way ANOVA with Tukey’s multiple comparison. **** p-value <0.0001, *** p-value <0.0007. **F)** Violin plot displaying log-normalised *C5AR1* expression between cMos, preOC-1, and preOC-2 clusters as point of comparison. **G)** Violin plot displaying log-normalised *NFATC1* expression between clusters, split by disease state (PsA red, healthy cyan). Annotated clusters indicted on left of plot. **H)** Dotplot heatmap displaying log-normalised gene expression of key marker genes across the observed clusters. Dot size indicates proportion of cells with nonzero expression. Relevant functional groups indicated to left of plot. Pathways shown are those related to osteoclastogenesis and function. **I)** Heatmap of top 20 differentially expressed inferred transcription factor activities (regulon scores) across the annotated clusters, calculated using SCENIC analysis^142^. Number of genes associated with each regulon indicated in parentheses. **J)** Heatmap showing transcriptomic similarity (mean prediction score) of single cells within VASA-seq scRNAseq dataset, using *in vitro* generated osteoclastogenesis atlas as a reference. Scores were calculated using FindTransferAnchors in Seurat. **K)** Quantification of single-cell enrichment scores for late preOC signature from the *in vitro* atlas across VASA-seq clusters. P values calculated using two-tailed Mann-Whitney-U. ****p≤0.0001.

To further investigate preOC identity, we utilised SCENIC to infer transcription factor (regulon) activities within each cluster (**Figure 4I**). Classical monocytes are enriched for *MECOM*, a known target of *PU.1, miR-22*, and *CEBPA* during monocyte-to-macrophage differentiation^36^, as well as *STAT5b*, a regulator of GM-CSF signalling during monocyte differentiation^37^. PreOC show enrichment for the known osteoclastogenesis transcription factors *NF-κβ, JUN, FOS, REL*, *STAT3*, and *PRDM1* (also known as *BLIMP1*)^38–41^ (**Figure 4I).**

Finally, we compared the identity of our preOC to *in vitro* generated monocyte-osteoclast intermediaries. We used two publicly-available datasets^10, 42^ to generate an reference dataset (**Supplementary Figure 4B**) after removal of contaminating cells from the bone marrow derived datasets (neutrophils, B cells, mesenchymal stromal cells, *VCAM1*^pos^ stromal macrophages). Final populations are annotated based on key canonical markers, differentially expressed genes, and the source publications. Five key clusters are identified, monocyte-derived macrophages, early preOC, late preOC, proliferating preOC, and mature osteoclasts (**Supplementary Figure 4C**). Monocyte-derived macrophages are mostly found at d0 in the murine object, and featured genes common to cMos and monocyte-derived macrophages (*S100A8, CCL2, C5AR1, CCR2*) (**Supplementary Figure 4D-E**). Mature osteoclasts form a clear separate tail from the main clusters, are found in significant numbers after 3d of culture of murine bone marrow-derived macrophages (**Supplementary Figure 4D**), and feature expression of known key osteoclastic markers (*ACP5, CTSK, MMP9, ATP6V0D2, FOSL2, NFATC1*) (**Supplementary Figure 4E**). Generally, both early and preOC did not feature distinct gene expression profiles but progressively downregulate mono-mac genes and upregulate osteoclast-related genes over the course of their differentiation (**Supplementary Figure 4E**). Finally, a population of proliferating preOC is identified, featuring low expression of mono-mac genes, similar levels of osteoclastic genes to early preOC, and high expression of markers of proliferation (*MKI67, TOP2A, BIRC5*) (**Supplementary Figure 4E**).

By using reference-based label transfer, we analysed the transcriptomic similarities between our assigned clusters and the final clusters identified from the *in vitro* generated osteoclast reference dataset (Error! Reference source not found.**Figure 4J-K**). Each of our annotated plate populations (cMos, preOC-1, preOC-2) display similarities to the corresponding populations (mono-macs, early preOC, late preOC) from *the in vitro* generated osteoclastogenesis reference dataset (**Figure 4J**). The enrichment for the late preOC signature is progressively upregulated between clusters, supporting the notion of transitioning cells (**Figure 4K**). In summary, our findings confirm that our sorted populations represented *bona fide* preOC and suggest a differentiation pathway from cMos to preOC-2.

### Classical monocytes form preOC *in silico*

It has been widely hypothesised that specific monocytic precursors are committed to differentiation towards an osteoclast fate^9, 12, 13, 43–52^. Therefore, we more formally investigated the existence of a potential trajectory, as well as changes in gene expression along this. We performed trajectory analysis by using Slingshot^53^ (**Figure 5A**). This identifies a clear trajectory from cMos to preOC-2 in keeping with this hypothesis. Inferred trajectories can be subject to a variety of limitations including requiring users to designate the starting cells, overfitting, and being unable to predict transcriptomic states before or after an initial state^54^. We therefore performed scTour, a deep learning-based architecture that overcomes many of these limitations^54^, on our dataset. This demonstrates a similar trajectory of cells from cMos to preOC-2 (**Supplementary Figure 5A-B**), further supporting the existence of this trajectory.

**Figure 5.**
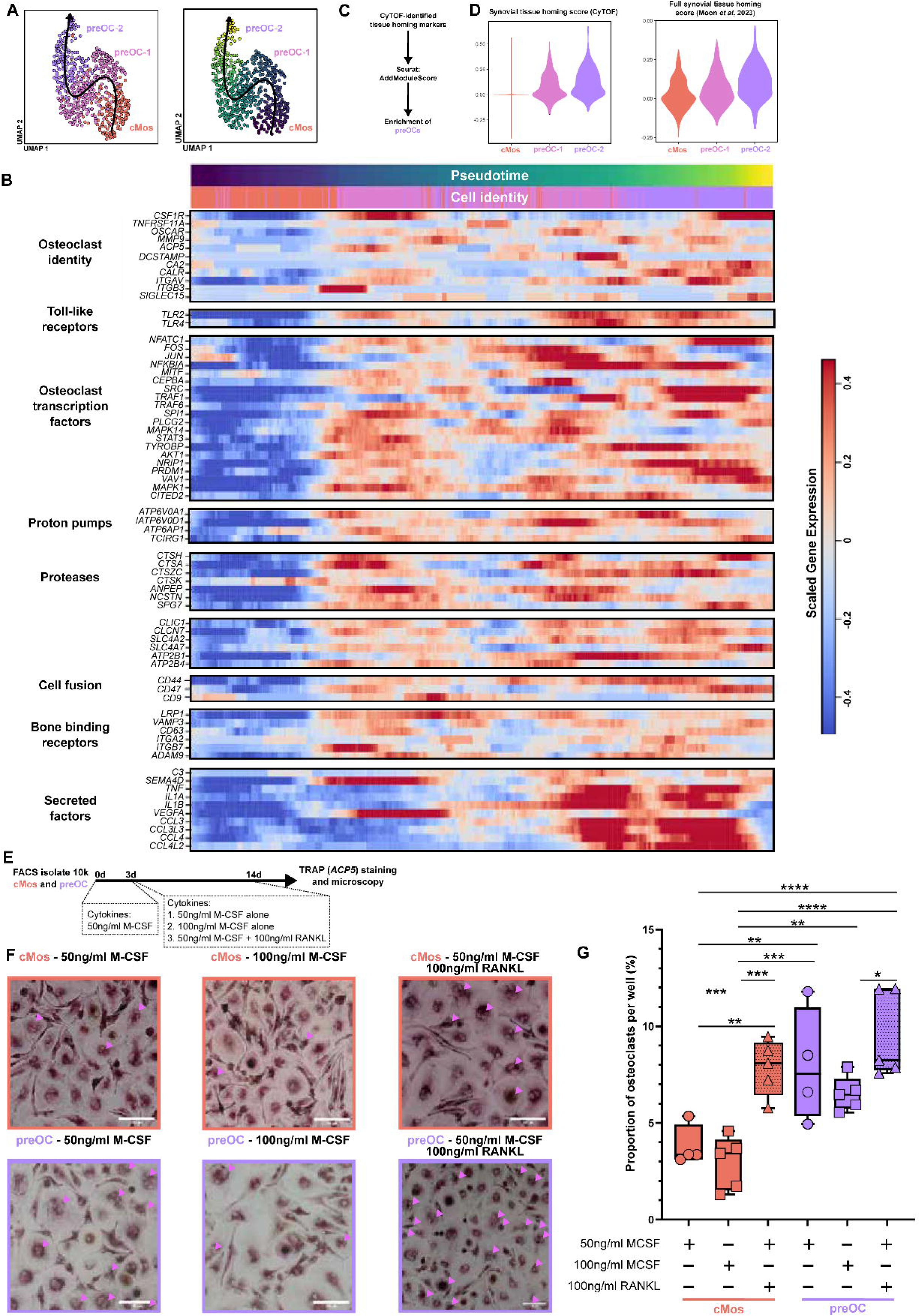
Pre-osteoclasts are primed for tissue homing and form osteoclasts in vitro in a RANKL-independent manner. **A)** Left panel; UMAP visualisation of 658 classical monocyte and preOC single cell transcriptomes with Slingshot trajectory overlaid (black arrow). Right panel; UMAP visualisation of cMos-preOC transcriptomes with cells coloured by Slingshot pseudotime (dark blue to yellow) and arrow overlaid. **B)** Heatmap of smoothed normalised gene expression of select osteoclast relevant genes over Slingshot pseudotime. Process relevant to displayed genes displayed to left. Annotation bars (top) show position within pseudotime (dark blue to yellow) and by cell identity as per base UMAP visualisation. **C)** Schematic used to score VASA-seq clusters for relevant markers of tissue homing. **D)** Enrichment for suspension mass cytometry identified markers on plate-based populations. Left panel, violin plot showing expression of AddModuleScore of CyTOF-tissue homing scores. Right panel, violin plot showing scores of cMos, preOC-1, and preOC-2 for the full synovial tissue homing score described by Moon et al, 2023^231^. P-value calculated by 2-way ANOVA with Tukey’s multiple comparison. ** p-value = 0.001, **** p-value <0.0001. **E)** Schematic depicting the experimental design for osteoclastogenesis assay. **F)** Brightfield microscopy images of cultured cMos (pink, top panels) or preOC (purple, lower panels) at d14 after staining for tartrate-resistant acid phosphatase. Relevant culture conditions described above each brightfield image. Panels show representative output at higher magnification. Pink arrows indicate TRAP^pos^ multinucleated cells. Scale bar, depicted in white is 250µm **G)** Boxplot demonstrating the proportion of osteoclasts (TRAP^pos^, ≥3 nuclei) produced per well. Each dot represents one biological replicate (N=5). Boxes and dots are coloured by sorted identity (cMos pink, preOC purple). P-values calculated using two-way ANOVA with Dunnett’s multiple comparison test. *p-value≤0.05, ** p-value≤0.01, *** p-value≤0.001, **** p-value≤0.0001.

The expression of genes of interest was profiled over Slingshot pseudotime. These analyses show that the cells upregulated markers relevant to osteoclastogenesis over pseudotime (**Figure 5B**). This includes known markers of osteoclast identity (*DCSTAMP, ACP5, CA2, CALR, ITGAV, SIGLEC15*)^8, 10, 29, 31, 42, 55, 56^, transcription factors including *NFATC1*, as well as the machinery required for bony resorption (*ATP6V0D1, TCIRG1, CTSK, ANPEP*)^12^. Components important for cell: cell fusion (*CD44, CD47, CD9*)^14, 29, 31^ and pro-inflammatory cytokines (*CCL3, CCL4, TNF, IL1α*) increase over differentiation (**Figure 5B**). Additionally, *SEMA4D*, a marker expressed by osteoclasts and that acts as a mediator of osteoclast-osteoblast interactions^57^ is upregulated over pseudotime. These findings indicate that we have successfully captured cMos committed to osteoclast cell fate.

### Psoriatic preOC display transcriptomic differences to healthy preOC

We next determined if disease state has any effect on preOC. We identified DEGs between healthy preOC-1 and PsA preOC-1, and healthy preOC-2 and PsA preOC-2. Minimal (10) DEGs are found between healthy and PsA preOC-1 (**Supplementary Figure 4F**). A relevant marker upregulated in disease state is TLR2, an important pattern-recognition receptor for osteoclasts, and important in binding to SIGLEC15 to initiate fusion of preOC^58, 59^. There are 203 DEGs between PsA preOC-2 and healthy preOC-2, with 149 upregulated in PsA preOC-2 (**Supplementary Figure 4G**). This includes upregulation of *CCL4, NTN1, SEPTIN9, IRAK2*, and *ATP6V1C1*, genes that are important for the migration, differentiation and function of preOC / mature osteoclasts^59–63^. These data indicate that psoriatic preOC-2 are more primed for osteoclastogenesis fate than their healthy counterparts.

### PreOC are primed for a synovial tissue homing signal

A key feature of the preOC population identified by the suspension mass cytometry profile is enrichment for markers of synovial tissue homing. We therefore looked for similar enrichment within our sorted populations, to further support their identity and potential role in psoriatic pathophysiology. We assessed for enrichment of both the full synovial tissue homing score^33^ and the profile we identified using suspension mass cytometry (**Figure 5C).** In keeping with our prior findings, the preOC populations feature significant upregulation of the CyTOF-identified preOC tissue homing signature (**Figure 5D)**. The full synovial tissue homing signature detailed in Moon *et al*^33^ is also significantly enriched along the cMos to preOC-2 differentiation (**Figure 5D)**.

### PreOC form osteoclasts *in vitro* in a RANKL-independent manner

We next sought to confirm that the preOC are capable of forming osteoclasts *in vitro*. We purified cMos or preOC from healthy donors by FACS (**Supplementary Figure 5C**) and plated them at a density of 10,000 cells per well. Cells were supplemented with cytokines that promoted either standard monocyte-derived macrophage differentiation, or osteoclast formation^8^. Cells were cultured for 14d and stained for the osteoclast marker tartrate resistant acid phosphatase (TRAP), with osteoclasts defined as TRAP^pos^ cells featuring at least 3 nuclei in keeping with standard microscopy definitions^6–8^ (**Figure 5E**).

We find osteoclast formation is rare when cMos cells are stimulated with M-CSF (50ng/ml and 100ng/ml alone) (**Figure 5F, G, Supplementary Figure 5D**). Addition of 100ng/ml RANKL produces osteoclasts (**Figure 5G**). In comparison, sorted preOC produce osteoclasts upon supplementation with exogenous RANKL (**Figure 5G**). However, they also produce osteoclasts in a RANKL-independent manner, for both cells cultured with 50ng/ml or 100ng/ml M-CSF alone (**Figure 5G**). There is a small reduction in osteoclast numbers from sorted preOCs when they were cultured with 100ng/ml M-CSF alone in comparison to preOC cultured with RANKL (p-value<0.05). This is in keeping with reports that higher concentrations of M-CSF can inhibit osteoclastogenesis^64^. In conclusion, our findings demonstrate that preOC are biased towards osteoclast fate in comparison with cMos and can produce osteoclast without exogenous RANKL.

### PreOCs and osteoclast-lineage cells are found in synovial tissue at single-cell resolution

We next determined if preOC and osteoclast cells were expanded in psoriatic synovial tissue. We obtained our own scRNA-seq data from healthy and active treatment naïve PsA synovial samples, and integrated this with 6 publicly available synovial datasets from RA, healthy controls, osteoarthritis (OA) and undifferentiated inflammatory arthritis (UA)^65–70^ (**Figure 6A**) to produce a reference atlas of 638,506 cells (156 donors) in synovial tissue (**Figure 6B, Supplementary Figure 6A-B**). We first annotated broad cell classes (myeloid, stromal cells, endothelial cells, T cells, B / plasma cells, pericytes, NK cells, and mast cells) for all donors and disease states (**Figure 6B-C, Supplementary Figure 6C**). These lineages were annotated based on cell type harmonisation across datasets using CellHint^71^. We reanalysed the myeloid subset of the data and utilised the gene signature generated from our plate-based data to identify preOC (**Figure 6D, Supplementary Figure 6D**). Other subpopulations were annotated using Alivernini *et al*, 2020 and Zhang *et al*, 2024 as the references (**Figure 6E**). We found considerable overlap between the MERTK/S100A8 macrophages from the Zhang dataset^67^ with the TREM2^high^ and TREM2^low^ subsets found in the Alivernini dataset^65^. The initial cluster 10 most closely resembled the TREM2^high^ cells, and cluster 4 demonstrated highest enrichment for MERTK/S100A8 macrophages though sharing considerable enrichment also for TREM2^low^ and TREM2^high^ cells. We identified osteoclast-lineage cells (OCL) on the basis of their expression of key canonical markers of mature osteoclasts (**Figure 6D, F**). However, due to the limitations of scRNA-seq, we were not able to demonstrate they are mature multinucleated cells, hence we termed them OCL-1 and −2 (**Figure 6E, F**). Within each other population, sub-populations were annotated using the identities from Zhang *et al*, 2023^67^ (**Supplementary Figure 6E-I**). The presence of OCL cells in synovial tissue suggests that infiltrating preOC undergo further differentiation into mature cells *in situ*. We performed trajectory analysis using scTour (**Figure 6G**) and other methods (**Supplementary Figure 6J**) on preOC and osteoclast-like subsets, revealing a trajectory from preOC to OCL-2. PreOC express key transcriptional markers expressed in the plate-based data including *ELMO1* and *DOCK4*, and osteoclast-lineage cells expressed osteoclast markers including *CTSK, ACP5*, and *MMP3* (**Figure 6H**). OCL-1 and OCL-2 expressed similar markers, with OCL-2 also expressing known osteoclast-derived coupling factors (*CST3*)^72^, and other markers of osteoclast maturity (*MMP3*^73^*, CLU*^74^) (**Figure 6H**).

**Figure 6.**
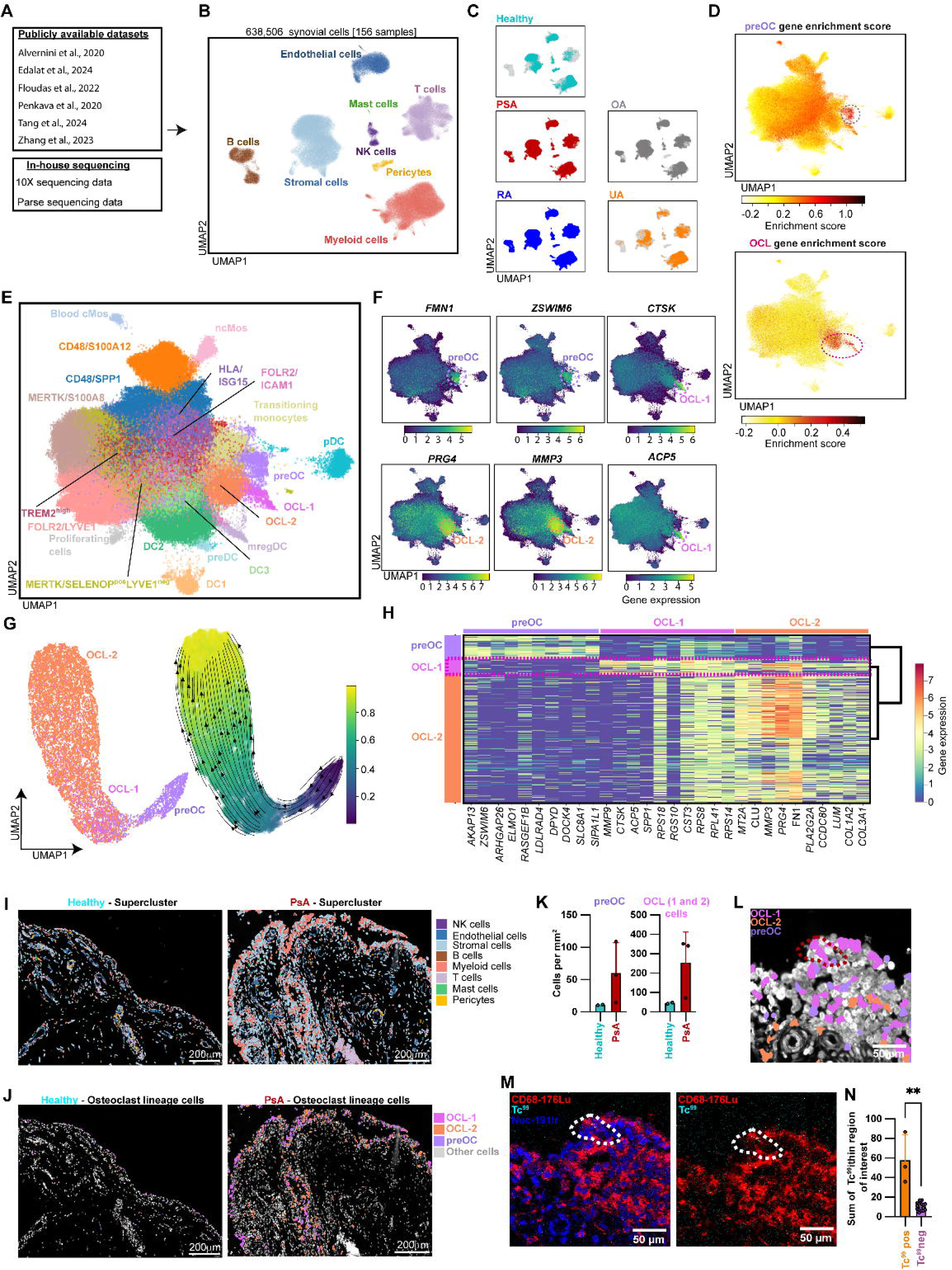
Pre-osteoclasts are enriched in synovial tissue from active psoriatic arthritis and are primed for migration into tissue. **A)** Datasets used to formulate Human Synovial Tissue Atlas. **B)** UMAP visualisation of 638,506 synovial cells from all datasets (156 donors), coloured by supercluster. **C)** UMAP visualisation of cells coloured by disease state. **D)** UMAP visualisation of myeloid subclustering, coloured by gene enrichment score for VASA-seq derived preOC gene signature (top). UMAP visualisation of myeloid subclustering, coloured by gene enrichment score for OCL gene signature captured from a bulk RNA-seq experiment (bottom)^41^ **E)** UMAP visualisation of myeloid subclusters with final assigned annotations. **F)** UMAP visualisations of key markers of interest on E, coloured by gene expression. Regions of interest for preOC, OCL-1 and OCL-2 indicated by coloured dotted circles. **G)** Left panel, UMAP visualisation of subsetted preOC, OCL-1 and OCL-2 cells. Right panel; trajectory analysis of subsetted preOC, OCL-1, and OCL-2 as elucidated by scTour. UMAP visualisation is coloured by pseudotime, with arrows indicating direction. **H)** Heatmap demonstrating key markers of interest across the preOC, OCL-1, and OCL-2 populations. **I)** Representative view of Xenium spatial sequencing of healthy and PsA synovium. Cells are coloured according to their supercluster identity. Scale bar, depicted in white is 200µm. **J)** Representative view of Xenium spatial sequencing of healthy and PsA synovium. Cells of osteoclast lineage are coloured according to their identity. Scale bar, depicted in white is 200µm. **K)** Boxplot demonstrating the number of preOC and osteoclast-lineage cells (1 and 2) per mm2 of synovial tissue. Each dot represents one biological replicate (N=5, 2 healthy and 3 PsA). Statistical analysis not possible due to limited sample numbers. **L)** Representative view of the Xenium spatial sequencing from MONOQUANT2. Region of interest enriched for Tc^99^ signal indicated with red dotted line. Cells are coloured according to identity as preOC, OCL-1, and OCL-2. **M)** Representative view of the imaging mass cytometry performed after Xenium workflow from MONOQUANT2. Region of interest enriched for Tc^99^ signal indicated with white dotted line. Marker-conjugate shown illustrated in top left of panels. Scale bar, depicted in white is 50µm.Left panel; DNA-191Ir (blue pseudocolour), CD68-176Lu (red pseudocolour), and Tc^99^ (cyan pseudocolour) respectively. Right panel; CD68-176Lu (red pseudocolour), and Tc^99^ (cyan pseudocolour) only. **N)** Boxplot demonstrating the Tc^99^ signal found within positive cells and surrounding Tc^99-neg^ cells. Each dot represents one cell. P-values were calculated utilising Mann-Witney U, **p-value<0.01.

### PreOCs are expanded in psoriatic synovium as demonstrated via spatially resolved transcriptomics

We next sought to identify preOC and osteoclast lineage cells *in situ*. From psoriatic donors who had undergone Tc^99m–^monocyte imaging, ultrasound guided synovial biopsies were taken 24-hours after imaging. Healthy synovial tissue was obtained from near-healthy donors. Spatial transcriptomics was performed on N=3 active PsA and N=2 healthy donors using the 10x Xenium 5K human immune panel with 50 genes added on. The spatial data were annotated at the population level by using the synovial atlas as the reference (**Figure 6I, Supplementary Figure 7**). Focusing on the myeloid group, we annotated sub-populations in a similar fashion, with preOC and OCLs identified using the top 100 genes from the plate-based data (**Figure 6J**). PsA tissue features enrichment for both preOC and osteoclast-lineage cells, in comparison to healthy tissue (**Figure 6K**). The MERTK^pos^ S100A8^pos^ population was also localised to the lining layer in health (**Supplementary Figure 7G, right panel**), but spatially disrupted in PsA (**Supplementary Figure 7G, left panel**) in agreement with the findings of others^65^.

### Previously radiolabelled cells are detectible *in vitro* using suspension and imaging mass cytometry

We hypothesised that technetium-99, as a heavy metal, would be detectable via suspension and imaging mass cytometry. We radiolabelled donor-derived monocytes *in vitro*, and froze these until they were completely decayed. Labelled and unlabelled monocytes were then thawed, and cultured for up to 14 days. Cells were removed from the plates at different time points, and stained using a simplified mass cytometry panel. An aliquot from the stained cell suspensions was transferred to glass slides via cytospin, and allowed to airdry. Cells were then assessed by both suspension and imaging mass cytometry (**Supplementary Figure 8A**), and the incorporation of Tc^99^ assessed (**Supplementary Figure 8B**). We found that it was possible to discriminate technetium^-99^ signal in labelled cells (**Supplementary Figure 8C**) via suspension mass cytometry. Signal was most prominent in cells at d0 in culture, with progressive loss of signal over 7 days (**Supplementary Figure 8C**). Signal was not readily detectable by 14d, though analysis in these cells was restricted by the limited number of monocyte-derived macrophages that survived to this timepoint, in both unlabelled and labelled donors (**Supplementary Figure 8C)**. With imaging mass cytometry, Tc^-99^ signal was not detected in unlabelled cells, though monocytes and macrophages could be clearly identified (**Supplementary Figure 8D**). Tc^99^ signal in labelled cells was rare, though occasionally detectible by imaging mass cytometry (**Supplementary Figure 8E**). Small foci of Tc^99^ signal were seen within cell boundaries, and tightly associated with both nuclei and myeloid signals (**Supplementary Figure 8E)**. These cells have a much higher density of technetium-99 within their boundaries than surrounding non-positive cells (**Supplementary Figure 8F**). This confirmed that it is possible to detect Tc^99^ labelled monocytes by both suspension and imaging mass cytometry.

### PreOC migrate to psoriatic joints

Finally, we determined if the tissue homing profile of preOC drives preferential recruitment to psoriatic joints. Immediately after the completion of the xenium workflow, slides were transferred to CyTOF-grade PBS and then imaging mass cytometry staining and imaging was completed. One of the two PsA donors had three foci of Tc^99^ positive cells within their synovial tissue, localised to the lining layer (**Figure 6L-M**). The other donor, and healthy donors, contained no detectable Tc^99^ positive cells. We superimposed the imaging mass cytometry staining over the gene expression data, demonstrating that the three foci of Tc^99^ positive cells all co-localised to cells of the osteoclast lineage. One representative example of this is shown in **Figure 6L-M**. These cells have a much higher density of technetium-99 within their boundaries than surrounding non-positive cells (**Figure 6N**). This supports our hypothesis that the homing profile found on preOC is important for migration into synovial tissue.

## Discussion

While it is established that monocyte-derived cells can generate mature osteoclasts with appropriate stimulation, the precise identity of circulating precursors has remained elusive. Early reports suggested that subsets of myeloid cells, including cells annotated as pre-dendritic cells^6^ or specific synovial macrophage populations^7^, can form osteoclasts *in vitro*. Efforts to define preOC phenotypes have produced evolving and sometimes conflicting marker sets, ranging from CD16^pos^ intermediate monocytes^8^ to CSF1R^high^ CX3CR1^high^ cells^9, 10^. A major barrier has been the technical difficulty of extracting and sequencing multinucleated osteoclasts *in vivo*^9^, leaving most insights reliant on *in vitro* systems^10, 42^. Murine studies have described both homeostatic and inflammatory preOCs, with the latter expanding during chronic inflammation, displaying enhanced migratory capacity, and producing highly resorptive osteoclasts^12^. However, the reported surface marker profiles are shared widely across monocytes^12^, raising the possibility as to whether these populations truly represent distinct committed precursors or rather reflect activation states imposed by the inflammatory milieu.

Our study provides a comprehensive characterisation of circulating human preOCs, integrating phenotypic, transcriptomic, and functional data. Using multiparameter mass cytometry and a refined flow-cytometry strategy, we were able to isolate preOCs. Transcriptomic analyses revealed close similarity of circulating preOC to *in vitro* generated monocyte–osteoclast intermediates, including progressive upregulation of osteoclastogenic genes and strong enrichment for *NFATC1*, the master regulator of osteoclast differentiation. Functionally, these preOCs form multinucleated osteoclasts, even without exogenous RANKL, distinguishing them from classical monocytes. The latter suggests that these cells are already primed for differentiation, potentially through prior exposure to RANKL^43, 75–77^ or non-canonical pathways such as TGFβ/TNF-mediated programming^78^, both relevant in PsA.

Comparison with prior descriptions places our population closest to CD14^pos^ CX3CR1^pos^ DCSTAMP^high^ cells, which had previously been noted to generate more osteoclasts than their DCSTAMP^low^ counterparts but were not extensively characterised^11, 13, 14^.

A striking feature of our preOCs was their expression of a suite of homing receptors, including CCR3, CCR4, CCR7, CCR9, CCR10, CXCR4, and CX3CR1, together with adhesion molecules such as CD44, EPCAM, and integrins (ITGA7, ITGA9, ITGB8). These profiles suggest specialised potential for recruitment to inflamed synovial tissue and bone. For example, CCR4 ligands (CCL2, CCL5^79^) and CXCL12–CXCR4 interactions are enriched in inflamed joints and known to promote osteoclastogenesis^80–83^. Similarly, ITGA9 binding partners such as tenascin-C and osteopontin are abundant in psoriatic synovium^84^. ITGA7 has also not been reported on preOC or mature osteoclasts previously, though its role in the recruitment of macrophages to sites of periosteal injury via tenascin has only been recently elucidated^85^. This indicates a potential role for the positioning of preOC to bone implicating this ITGA7/ITGA9-tenascin axis in pathological recruitment of preOCs to sites of bone erosion^84^. This also poses interesting questions for the role of preOC in pathological new bone formation at sites of periosteal/entheseal inflammation, given the tight coupling of resorptive activity with osteoblast activity in bone remodelling^86^.

Notably, we observed *NFATC1* upregulation even in classical monocytes from PsA donors, suggesting systemic priming of the osteoclastogenic programme in disease. This is in keeping with elevated RANKL levels in PsA serum and synovial fluid, as well as the central role of the key therapeutic targets TNF, IL-23, and IL-17 in driving pathology. Another possibility is priming of cells in psoriatic skin, as recently demonstrated for other myeloid precursors in psoriatic disease^87^.

The ability of preOCs from healthy donors to form osteoclasts without exogenous RANKL raises intriguing mechanistic questions. While autocrine RANKL production has not been demonstrated in preOCs, alternative pathways for osteoclastogenesis exist. TGFβ signalling can epigenetically programme myeloid cells to adopt an osteoclast fate in response to TNF, bypassing canonical RANKL-RANK interactions^78^. PsA is characterised by heightened TGFβ activity^88, 89^, but the presence of RANKL-independent osteoclastogenesis in healthy donors suggests that preOCs may sometimes receive sufficient priming signals in circulation to commit to this fate. This study also finds that final cell fate can be significantly biased even before a cell enters synovial tissue in humans.

Several limitations of our approach should be acknowledged. First, it is difficult to ascertain whether mature multinucleated osteoclasts are adequately represented in both our single-cell atlas and spatial datasets. scRNA-seq is known to incompletely capture large, adherent, multinucleated cells such as osteoclasts, which are technically challenging to dissociate and recover from tissue samples. In addition, previous studies indicate that osteoclasts are most frequently concentrated at the synovium–bare bone interface^7, 90^, a region that is not readily accessible through ultrasound-guided synovial biopsy in humans. This anatomical limitation may further reduce the likelihood of sampling fully mature osteoclasts. Nevertheless, we observe co-localisation of osteoclast-lineage cells within the thickened lining layer of psoriatic pannus. This spatial pattern is consistent with cellular fusion and multinucleation, suggesting the presence of cells undergoing the terminal stages of osteoclastogenesis.

Prior work using radiolabelled monocytes estimated that a median of 323 labelled cells trafficked to inflamed knees in patients with RA following autologous reinfusion^22^. Although our study was not specifically designed to quantify cellular influx in this manner, our findings are broadly consistent with these observations.

The two PsA tissue sections analysed had volumes of 47.5 mm³ (MONOQUANT1) and 69.25 mm³ (MONOQUANT2), representing less than 1% of even the lower estimates of total knee synovial tissue volume measured by MRI in OA^91^. This limited sampling volume may explain why, in MONOQUANT1, we detected monocyte accumulation in inflamed joints on imaging but did not identify any Tc^99^-^pos^ cells by IMC. The ability to trace and phenotype circulating cells as they enter tissues represents a critically needed advance for human immunology. We term this technique mass-assisted single- and spatial sequencing (MASS-seq).

In contrast, in the PsA donor MONOQUANT2, three Tc^99^-^pos^ foci were identified. All three exhibited a phenotype consistent with osteoclast-lineage cells. Although the possibility that Tc^99^ leached from monocytes present within the tissue cannot be completely excluded, this appears unlikely. Instead, we propose that these findings reflect preferential homing of preOCs to inflamed joint tissue compared with circulating monocytes, in support of our central hypothesis and broader data.

Tracing technetium-labelled cells using IMC presents technical challenges, particularly in the context of low labelling efficiency and the inherent limitations of analysing two-dimensional cross-sections through cells. These constraints may be mitigated by utilising “tissue mode” acquisition on the Hyperion system (5µm resolution, compared with the 1µm cell-focused mode), thereby increasing technetium ion density per pixel. Alternatively, optimisation of radiolabelling strategies, including the use of different ligands or methodologies, may further enhance detection sensitivity.

In conclusion, we define a distinct population of circulating human preOCs with a transcriptional programme, surface phenotype, and functional behaviour consistent with cells primed for osteoclastogenesis. Unlike conventional monocytes, these preOCs can differentiate into osteoclasts without exogenous RANKL, produce pro-inflammatory cytokines, and express homing receptors that may direct them to sites of pathological bone resorption. Their expansion and activation in PsA provide a mechanistic link between systemic inflammation and local bone destruction. Future studies should determine whether targeting preOC recruitment or priming offers new therapeutic avenues for inflammatory arthritis.

## Materials and methods

### Patient samples

All samples were obtained utilising written informed consent from participants under ethical approval obtained from the relevant research ethics committees as detailed below. Two separate ethics approvals were used for this thesis. All aspects were carried out in accordance with the guidelines under the Declaration of Helsinki 2000.

Work comprising the sampling and analysis of blood from healthy, PsA, and PsC donors and synovial tissue specimens from near-healthy and PsA donors was performed under approval 19/EE/0316 and integrated research application system (IRAS) 271612 from the Central Cambridge Research Ethics Committee.

Work comprising the isolation, labelling, and imaging of Tc^99m–^monocytes in healthy and active PsA (including biopsy for PsA donors) was performed under approval 22/EE/0228, IRAS 304702 from the East of England Essex Research Ethics Committee.

### Reagents used

4% chlorhexidine solution (Hibiscrub 4%, LCKW2), 2% Lidocaine (Hameln, LD1002), 10% neutral buffered formalin (CellPath BAF-6000-08A), RPMI media (Gibco, 31870-025), 10% foetal calf serum (FCS, Hi-FBS; Sigma-Aldrich, F9665-500ml), penicillin-streptomycin (Pen/Strep, Gibco, 15140-122), Amphotericin B (Sigma-Aldrich, A9528), Gentamycin (Sigma-Aldrich, G1272), L-glutamine (Thermofisher Scientific, 25030024), sterile 5% acid-citrate-dextrose (ACD, Huddersfield Pharmacy Special, 321219X), sterile 0.9% normal saline (Fresenius kabi, 232250), Pancoll (density 1.077 mg/ml; Pan-biotech, P04-60500), Trypan-blue (ThermoFisher, 15250061), CryoStor CS10 (STEMCELL Technologies, 100-1061), Liberase-TM (Thermolysin Medium, Sigma, 5401119001), xylene (Fisher Scientific, X/0250/17), Harris acidified haematoxylin (PFM Medical, PRC/R/51), 1% aqueous eosin Y (PFM Medical, PRC/66/1), DePeX (BDH, 361252B), Zombie-Aqua (Biolegend, 423101), 5% human AB (Sigma-Aldrich, H4522), 1% mouse (Sigma-Aldrich, M5905), 1% rat serum (Sigma-Aldrich, R9759-5ML), 2mM ethylenediaminetetraacetic (EDTA, Sigma-Aldrich, 102499418), 1X BD CellFix (BD Biosciences, 340181), DAPI (Sigma-Aldrich, D9542), Live-dead fixable blue dead cell stain kit (ThermoFisher, L23105), transcription factor buffer set (BD Biosciences, 562574), APC Lightning-Link kit (abcam, ab201807), UltraComp eBeads Plus Compensation Beads (Invitrogen, 01-3333-42), MaxPar X8 (lanthanide metals), MCP9 (cadmium metals), and cisplatin X8 (platinum) antibody labelling kits (Standard Biotools), TCEP (Sigma-Aldrich, 646547-10X1ML), 0.05% sodium aside (Boca Scientific, 131 050), DNase I (Roche, 10104159001), Cell-ID rhodium (Standard Biotools, 201103A), Maxpar Cell staining buffer (Standard Biotools, 201068), human TruStain FcX (Biolegend, 422301), 16% methanol free formaldehyde (ThermoScientific, 28906), 125nM Cell-ID Intercalator Iridium (Standard Biotools, 201192A), Maxpar cell acquisition solution (Standard Biotools, 201240), EQ normalisation beads (Standard Biotools, 201245), 1X target retrieval solution (Agilent, S2367), Maxpar water (Standard Biotools, 201069), 3% protease free bovine serum albumin (BSA) solution (Sigma-Aldrich, A3059-50G), Tween-20 (Sigma-Aldrich, P1379-100ml), CD14 MicroBead kit (Miltenyi, 130-050-201), MS column (Miltenyi, 130-042-201), LS column (Miltenyi, 130-042-202), M-CSF (Gibco, PHC9501), accutase (Biolegend, 423201), MEMα (ThermoFisher, 22571038), RANKL (Peprotech, 310-01C-10UG), TRAP staining kit (Sigma-Aldrich, 387A-1KT), RLT buffer (Qiagen, 79216), beta-mercaptoethanol (Sigma-Aldrich, M6250-100ml), scRNA purification kit (NORGEN, 51800), DNase-I (NORGEN, 25710), SMART-seq v.4 Ultra Low Input RNA Kit for Sequencing (Takara, R400752), RNase Zap (Invitrogen), EDTA collection tubes (BD, 367525), 6% hydroxylethyl starch (HES, Grifols, 0318), hetastarch (CFP-S), 30% (5M) sterile sodium chloride (South Devon Healthcare, 01883/6158R), water for injection (BP), sterile Percoll Plus (GE Healthcare, 17-5445-02), CliniMACS buffer (Miltenyi, 200-070-025), CliniMACS CD14 microbeads (Miltenyi, 170-076-705), sterile LS column (Miltenyi, 130-042-202), sterile hexamethylpropyleneamine oxime (HMPAO)

### Tissue Processing

#### PBMC

All samples were processed immediately upon receipt. Human blood samples were diluted 1:2 in phosphate buffered saline (PBS), layered onto a Pancoll gradient (PAN biotech, P04-60500) and centrifuged at 3000rpm for 20 minutes without brake. The PBMC interface was collected and washed in PBS. Platelets were depleted via centrifuging at 400*g* for 10 minutes without brake. Cells were resuspended then counted using a haemocytometer and Trypan-blue (ThermoFisher, 15250061). Cells were then either used for downstream applications or frozen at 5-50 million cells per ml in CryoStor CS10 (STEMCELL Technologies, 100-1061) in liquid nitrogen until use.

#### Synovium

All specimens were processed in line with the protocol described by Alivernini *et al*, 2020^65^. Briefly, human synovial samples were diced into 1mm^3^ fragments using sterile scalpels and transferred into a sterile universal container containing sterile RPMI with penicillin/streptomycin 100U/ml, L glutamine 2mM, and 0.15μg/ml Liberase-TM (Thermolysin Medium, Sigma, 5401119001). Tissue was incubated at 37°C for 30-45 minutes with agitation. After incubation, the mixture was passed through a 100μM cell strainer (Falcon, 352360) and residual cell clumps gently massaged using the sterile rubber end of a 1ml syringe plunger. Complete medium containing 10% FCS, antibiotics and L-glutamine as described above was poured through the filter to abort the digestion. The mixture was centrifuged at 400*g* for 10 minutes to pellet the cells. Cells were resuspended and counted using a haemocytometer. Finally, cells were frozen at 1-10 million cells per ml in Cryostor CS10 in liquid nitrogen until use.

#### Fixation of synovial tissues for microscopy / histology

All specimens for microscopy were fixed in 10% neutral buffered formalin for 24 hours immediately after they were obtained. After fixation, specimens were transferred into 70% ethanol and transferred to the histology service in the Department of Pathology for embedding in paraffin and sectioning. Briefly, specimens were sectioned at 3µm, deparaffinised in xylene (Fisher Scientific, X/0250/17), rehydrated in serial ethanol dilutions (100%, 95%, 70%), washed with tap water, and stained with Harris acidified haematoxylin (PFM Medical, PRC/R/51) for 10 minutes, and washed with alternating tap water and 1% acid alcohol made in house. Slides were then counterstained with 1% aqueous eosin Y (PFM Medical, PRC/66/1). Slides were then dehydrated in methylated spirits, cleared with xylene and mounted using DePeX (BDH, 361252B).

### Flow cytometry and sorting

#### Flow cytometry

Cells were blocked in human blocking buffer (5% human AB (Sigma-Aldrich, H4522) / 1% mouse (Sigma-Aldrich, M5905) / 1% rat serum (Sigma-Aldrich, R9759-5ML), 5% FCS, 2mM ethylenediaminetetraacetic (EDTA, Sigma-Aldrich, 102499418)) for 15 minutes at 4°C before being incubated with an antibody cocktail for 30 minutes at 4°C Antibodies used are listed in **Supplementary Table 5**. Cells were washed and resuspended in FACS buffer (PBS containing 2 % Hi-FBS and 2 mM EDTA). Cells were then washed and resuspend in 200μl of 1X BD CellFix (BD Biosciences, 340181) diluted 1:10 in double distilled water before analysis on Cytek Aurora with 5 lasers and 16 ultraviolet, 16 violet, 14 blue, 10 yellow-green, and 8 red detection modules (Cytek). The lineage (Lin) channel in flow cytometry analyses included combinations of the markers CD3, CD19, CD20, CD56, CD66b, CD235a and CD335, for the removal of contaminating T cells, B cells, NK cells and granulocytes.

#### Cell sorting

Cell suspensions for sorting were blocked, stained with the appropriate panel, and washed as described above except 5ml polypropylene round-bottom tubes (Falcon, 352235) were used. Cells were filtered through a 40μM filter (Falcon, 352340), pelleted then resuspended in 200-500μl of FACS buffer (PBS with 2% FCS and 2mM EDTA). Immediately prior to sorting, the suspension was stained for viability using 4’,6 diamidino-2-phenlyindole (DAPI, Sigma-Aldrich, D9542) at 1:1000 before sorted on the BD FACS Aria III using a 100μm nozzle into appropriate buffers. For the preOC sorts, cells were stained for 15 minutes at 37°C (for better binding of chemokine receptor markers^117^) and then 15 minutes at 4°C. Cells were then washed / treated as described above with the exception of an additional secondary staining. Primary staining included staining with an antibody against ITGAV conjugated to biotin. After washing off primary antibody cocktail, cells were incubated for 30 minutes at 4°C with streptavidin-BV785 and then washed. Immediately prior to sorting, the suspension was stained for viability using DAPI at 1:1000 before sorted using BD FACS Aria III using a 100μm nozzle or BD FACSDiscover S8.

### Suspension mass cytometry

Cells from liquid nitrogen were thawed in a water bath at 37°C and washed in pre-warmed RPMI media supplemented with 10% FCS and 40µg/ml DNase I (Roche, 10104159001). Thawed cells were washed twice in serum-free RPMI and DNase I and incubated at 37°C for 30 minutes. Cells were then filtered through a 100µM cell strainer, pelleted and resuspended in serum-free RPMI, at a maximum concentration of 10×10^6^ cells per 5ml polypropylene round-bottomed tube. Cells were incubated with 100µL of 1µM Cell-ID rhodium (Standard Biotools, 201103A) for 30 minutes at room temperature and then washed with 3ml of Maxpar Cell staining buffer (Standard Biotools, 201068). Cells were blocked using 10µL of human TruStain FcX (Biolegend, 422301) for 10 minutes at room temperature. Antibody cocktails were made up in Maxpar cell staining buffer in a final volume of 100µL. Antibodies used are detailed in **Supplementary Table 6**. Cells were stained in a water bath at 37°C for 10 minutes (to facilitate binding to chemokine receptors^117^) and then for a further 30 minutes at room temperature. Cells were washed with 2ml Maxpar cell staining buffer and fixed in 1.6% formaldehyde, diluted from 16% methanol free formaldehyde (ThermoScientific, 28906) in Maxpar PBS (Standard Biotools, 201058), for 10 minutes at room temperature. Cells were pelleted, supernatant removed, and stained overnight at 4°C with 1ml of 125nM Cell-ID Intercalator Iridium (Standard Biotools, 201192A), in Maxpar Fix and Perm Solution (Standard Biotools, 201067). Cells were then transferred to 1.5ml LoBind Eppendorf’s, with care being taken to collect residual cells. Cells were washed 1x in Maxpar cell staining buffer followed by 2x washes in Maxpar cell acquisition solution (Standard Biotools, 201240). Cells were counted and resuspended to a maximum of 1×10^6^ cells per ml in the 1.5ml Eppendorf. Immediately prior to acquisition, cells were admixed with EQ normalisation beads (Standard Biotools, 201245) and acquired on the Helios mass cytometer (Standard Biotools).

Samples were gated on manually in FlowJo (Treestar Inc) as described in the relevant results sections. For mass cytometry of PBMC specimens as part of the tissue homing workflow, cells were gated to the stage of live CD45^pos^ cells. UMAP was performed on each sample using the FlowJo plugin with the following parameters: CD3, CD19, CD20, CD66b, NKp46, HLA-DR, CD14, CD16, CD88, CD89, CD123, CD5, AXL, CD1c, FcER1A, CD163. This was to enable accurate gating out of a contaminating population of basophils (HLADR^low^, CD88^pos^, FcER1A^pos^, CD123^pos^, CCR3^high^) which otherwise are not easily separable from the monocyte gate using manual gating. Purified populations of MPs were then downsampled to 10k cells and exported as FCS files. These were imported in RStudio for more detailed analysis and loaded using the “cytof_exprsMerge” function from Cytofkit2^144^. Samples were arcsinh transformed using the “transform_asinh” function with cofactor 5 (as appropriate for mass cytometry data) from the R scales package^145^ and relevant metadata added. Samples were then loaded as Seurat objects from their dataframes, merged and scaled. Batch correction was performed on individual sample using the FastMNN package in SeuratWrappers^146^. Dimensionality reduction and clustering was performed using the “RunUMAP”, “FindNeighbours” and “FindClusters” commands using 44 PCs. Populations were annotated based on their expression of known canonical marker proteins. Differentially expressed markers were visualised using heatmaps, violin plots, and FeaturePlots.

The combined UMAP from above was exported as an FCS file and loaded into FlowJo. A gate was manually drawn around the putative preOC population, and the HyperFinder plugin in FlowJo was used to identify a gating strategy using default parameters.

### Imaging mass cytometry

#### Cytospins of radiolabelled cells

Cells were stained for mass cytometry as described above. Cells were resuspended at 0.5 million cells per ml in Maxpar cell acquisition solution. Cells were centrifuged onto glass slides using a Shandon Cytospin 2 (Marshall Scientific) at 150rpm for 3 minutes. Slides were placed into a slide box and allowed to air-dry before being analysed on the Hyperion XTI.

#### Post-xenium

After the xenium (10X) run on synovial tissue sections, slides were immediately placed within Maxpar PBS. The same process was repeated as above with the following alterations: the procedure began from incubating the slides with target retrieval solution as they had already undergone dewaxing and rehydration as part of the xenium workflow and the slides were not re-permeabilised with Tween-20 (as permeabilisation had been completed as part of the xenium workflow). Technetium-^99^ was included within the panel setup using the ruthenium^99^ channel as before. Antibodies used are listed in **Supplementary Table 6**. Samples when then analysed on the Hyperion XTI.

### In vitro culture of monocyte-derived macrophages

After PBMC isolation as described above, monocyte enrichment was carried out using positive selection with the CD14 MicroBead kit (Miltenyi, 130-050-201) as per the manufacturer’s instructions. Cells were blocked with FACS buffer at 80µL per 10 million cells, and labelled with CD14 MicroBeads at 20µL per 10 million cells for 15minutes at 4°C. Cells were washed with FACS buffer to remove unbound beads, and centrifuged at 400*g* for 5 minutes at 4°C. Cells were resuspended in 500µL of FACS per 100 million cells. A suitably sized MACS separator and column were assembled. An MS column (Miltenyi, 130-042-201) was used for ≤10^7^ cells and an LS column (Miltenyi, 130-042-202) used for ≥10^7^ but ≤10^8^ cells. Columns were prepared with FACS buffer, and the cell suspension washed through. Unlabelled cells were washed through with 3 washes through the column. 1ml of FACS buffer for MS columns or 5ml for LS columns was added, the column quickly removed from the magnet, placed over a 15ml Falcon tube and cells plunged from column into tube. The eluted CD14^pos^ cells were counted, and plated in RPMI containing 10% FCS, 20mM L-glutamine, and antibiotics at 1×10^6^ cells per ml on an appropriately sized sterile flat-bottomed plate, depending on the experiment as described below. Media was supplemented with M-CSF (Gibco, PHC9501) at 100ng/ml, and media refreshed every 3 days.

Cells were harvested by removing media from wells, and pre-warmed accutase (Biolegend, 423201) added to the relevant wells. This was incubated for 5-10 minutes at 37°C to loosen the cells, tapped vigorously, and the cells pipetted off and into FACS buffer. Wells were further washed 5x with FACS buffer. The removal of cells was checked using a light microscopy (EVOS XL Core, Life Technologies). The cells were washed twice in FACS before use in downstream applications.

### Osteoclast differentiation assays

10,000 cMos or preOC per treatment condition were sorted into minimal essential media alpha (MEMα, ThermoFisher, 22571038) supplemented with 10% FCS, L-glutamine, and antibiotics. Cells were counted using Trypan blue and haemocytometer, before being centrifuged and resuspended at 1000 cells per µL in MEMα supplemented with cytokines as described shortly. 60µL of MEMα as described below with relevant cytokines was pipetted into an 18-well Ibidi µ-slides (Ibidi, 81816). 10µL (equivalent to 10,000 cells) of cell suspension was then carefully pipetted into one corner of the plate, to a total volume of 70µL per well. Wells surrounding sample wells contained 100µL of PBS to reduce evaporation. Cells were cultured for 14 days before fixation and staining. Three conditions were plated:

1. 50ng/ml M-CSF for 14d
2. 50ng/ml M-CSF for 3d, followed by 50ng/ml M-MCSF and 100ng/ml RANKL (Peprotech, 310-01C-10UG)
3. 100ng/ml M-CSF for 14d.

At 14 days of culture, cells were fixed with 4% PFA for 20 minutes at 37°C, before being washed 3x in warmed deionised water. Cells were stained using the TRAP staining kit (Sigma-Aldrich, 387A-1KT). Briefly, TRAP staining solution was made by mixing warmed deionised water, diazotised fast garnet GBC base solution, naphthol AS-BI phosphate solution, acetate solution, and tartrate solution in the proportion recommended by the manufacturer. Cells were incubated with 100µL of this solution for 60 minutes at 37°C. Cells were washed 3x in warmed deionised water, before excess water was removed and the dish allowed to air dry in the dark. Wells were imaged under a 10X objective on a Zeiss AxioObserver Z1 microscope (Zeiss) using the tile scanning programme. Images were imported into Fiji^122^ for analysis. Briefly, 5 regions of interest of uniform size and placement were drawn over each of the wells, covering the majority of the well. Cells were manually counted in Fiji using its CellCounter plugin. The total number of cells, number of osteoclasts (defined as TRAP^pos^ cells containing at least 3 nuclei), and number of nuclei per osteoclast was enumerated.

### NanoZoomer slide scans of synovial sections

5µm sections of tissue adjacent to each section used for xenium sequencing were taken, and H+E stained by the histology team in the Cancer Research UK, University of Cambridge building, in the same way as described above. Finished slides were taken to the NanoZoomer S60 Digital slide scanner (Hamamatsu) in the Department of Pathology, and imaged. These were loaded in Fiji and exported as TIFFs.

### Bulk RNA-seq

Bulk-RNA sequencing of circulating blood MPS subsets of N=5 healthy and N=5 active treatment naïve PsA was performed. Briefly, a target of at least 500 cells were sorted into 600μl RLT buffer (Qiagen, 79216) containing 1% beta-mercaptoethanol (Sigma-Aldrich, M6250-100ml) within PCR-grade low-bind Eppendorf’s as described above using an Aria III. Upon sorting, the cells were vortexed for 60seconds to ensure cell lysis. RNA was precipitated by mixing with 600μl 70% ethanol and isolated using a scRNA purification kit (NORGEN, 51800) with DNA removal step using DNase-I (NORGEN, 25710). RNA was purified and frozen at −80°C until used. Quality-control of isolated RNA was performed by the Cambridge Stem Cell Institute Genomics Facility using the Eukaryote Total RNA Pico Chip (Agilent, 5067-1513). The same institution performed library construction and sequencing. High-quality RNA was used to construct Illumina mRNA sequencing libraries. cDNA synthesis and amplification were performed using a SMART-seq v.4 Ultra Low Input RNA Kit for Sequencing (Takara, R400752). The pooled libraries were sequenced using a NovoSeq S4 chip with 100 pair-end reads to yield an average of 90 million reads per sample. Reads were aligned with the Nextflow RNAseq pipeline (version 3.3) utilising the salmon_star option and returned as matrices by Russell Hamilton and Simon Andrews.

### Bulk-seq data analysis

After initial quality control, one healthy sample (HV002) was removed from the analysis due to high GC bias as discussed in Chapter 3. For the final analyses, the DC2 and CD14^neg^ CD163^neg^ DC populations were grouped together as these cells were transcriptionally similar and likely both represent DC2 as discussed in more recent publications^21^. Principle component analysis was performed in R^123^ using DESeq2^124^ with cell_type and Disease_state in the model. For cell:cell relationships in principal component (PC) space, the DESeq2 object was transformed using the varianceStabilizingTransformation function and visualised using ggplot2^125^. Hierarchical cluster analysis was performed using the “hclust” function using complete agglomeration as part of the “stats” package included as part of base R. The top 50 differentially expressed genes per MPS subset from the scRNAseq object were produced using the “FindMarkers” function in Seurat^126^ with the cluster of interest set as cluster 0, and with “NULL” as the comparator, before being collated into a gene list. The bulk-seq object was then log transformed and subsetted to this genelist and data plotted using pheatmap. Results for cell type comparisons between health and disease were extracted using the “lfcShrink” function, with the cell type comparison specified using the contrast parameter. Significantly differentially expressed genes (DEGs) were selected for each comparison with a LogFC > 2 and p_val_adj<0.05 and visualised using pheatmap. For pathway enrichment within cell types, normalised counts for both health and PsA cell types were extracted and combined as .txt files and loaded in Gene Set Enrichment Analysis (GSEA) application (Broad Institute)^127, 128^, with health vs. PsA set as the test phenotypes. Human hallmark, biocarta, keg, reactome, and wikipathways gene matrices were downloaded from the molecular signatures database (MSigDB)^127^ and analyses run with default parameters. Enriched pathways were manually explored and visualised. For tissue-homing markers, markers identified by differential gene expression and systematic visualisation of known tissue homing and adhesion markers were explored using manual lists and pheatmap^129^.

### In vitro generated osteoclast datasets

Data from scRNAseq of murine and human *in vitro* osteoclastogenesis were obtained as described above^50, 131^. Downstream analysis of each dataset was performed using Seurat (v4.0). The murine data from both datasets was filtered and humanised to genes containing only one-2-one human orthologues from ensembl^137^. For the murine datasets, cells with fewer than 750 genes, over 7000 genes or more than 5% mitochondrial genes were removed as per the base publications^50, 131^. For the human data, cells with fewer than 1000 genes or more than 10% mitochondrial genes were removed as per the authors workflow^50, 131^. Samples were merged, log-normalised and integrated by dataset following the Seurat v4 Integration workflow and 5000 variable features / integration anchors. UMAP dimensionality reduction was performed as described above using the first 30 PCs. Clusters were identified as described above. Significantly DEGs were identified using the “FindMarkers” function, using the Wilcox rank sum test, corrected for multiple comparisons. Clusters were identified in each dataset using the “FindNeighbours” and “FindClusters” functions in Seurat. Clusters were annotated based on the expression of known marker genes. Contaminating B cells, GMPs/neutrophils, and mesenchymal stromal cells were removed on the basis of markers from the base publications. After removal, the merged dataset was re-scaled, integrated, and analysed as described above. Final annotations were given based on key canonical markers reported in the source publications^50, 131^.

### Plate-based sequencing with vast transcriptome analysis of single cells by dA-tailing (VASA-seq)

Live single-cells of interest were isolated by FACS as detailed above and sorted into 384-well plates provided by Single Cell Discoveries. 8 empty well controls per plate are kept free of cells. Once cells were collected, plates were sealed, spun at 2000*g* for 2 minutes and snap-frozen using dry ice before storage at −80°C. Plates were then shipped to Single Cell Discoveries, Utrecht, Netherlands on dry ice. Sequencing was performed by Single Cell Discoveries to a target of 150,000 reads per cell on NovoSeq X Plus (Illumina). Read alignment and generation of count matrices from raw data were performed using STAR (v 2.7.11a) pipeline against the Human Genome (GRCh38-3.0.0) and performed unique molecular identifier (UMI) counting.

### VASA-seq data analysis

Initial preprocessing was completed by Single Cell Discoveries. FASTQ files were processed using Cutadapt^133^ to remove homopolymers and adaptor sequences. Ribosomal RNA was then removed using the Ribodetector^134^ package to filter out these unwanted reads. Pre-processed reads were then aligned to *Homo sapiens* (GRCh38) reference genome using STARsolo^135^. The count tables produced from these were then imported into Seurat for downstream analysis. Seurat data was provided as an RDS object. Downstream analysis of gene expression matrices was performed using Seurat (v4.0). Plates were loaded individually and relevant metadata and index sort data attached. Known empty control wells were removed. Cells with fewer than 500 genes or more than 10% mitochondrial genes were removed, yielding a total of 294 cells for plate 1, 223 cells for plate 2, and 219 cells for plate 3. Plate 1 had 4 wells containing doublets (identified at time of sorting), leaving a total of 725 cells (287 plate 1, 221 plate 2, 217 plate 3). Samples were merged, log-normalised and integrated following the Seurat v4 Integration workflow^126^ and 5000 variable features / integration anchors. UMAP dimensionality reduction was performed as described above using the first 15 PCs. Clusters were identified as described above. Significantly DEGs were identified using the “FindMarkers” function, using the Wilcox rank sum test, corrected for multiple comparisons. Clusters were identified in each dataset using the “FindNeighbours” and “FindClusters” functions in Seurat. Clusters were annotated on the basis of expression of known marker genes. For the purposes of this study 67 contaminating B cells (*CD79A^pos^*, *MS4A1^pos^*) were removed. After removal, the merged dataset was re-scaled, integrated, and analysed as described above. Volcano plots were visualised using the EnhancedVolcano^136^ package in R.

This and the *in vitro* osteoclast annotated datasets were loaded in R and the “FindTransferAnchors” command used within Seurat^126^, with the plate-based preOC data as the query dataset and the *in vitro* generated osteoclastogenesis dataset as the reference. 30 PCs were used, and the predicted identity added to the base preOC dataset in the metadata. Mean values for populations were calculated in R, and plotted with pheatmap^129^. Pseudotime trajectory analysis of cMos to preOC was performed using the Slingshot R package, with the start cluster specified^140^. The calculated trajectory was overlaid onto the UMAP embeddings. For the smoothed heatmap, genes of interest were taken and plotted along the trajectory using pheatmap^92^. A further confirmatory test of trajectory was performed using the scTour package^141^ using default setting, 3000 highly variable genes, and loss_mode of mean-squared error in python. SCENIC^142^ was used to infer transcription factor activity in our VASA-seq plate data. Analyses were performed in R. Briefly, regulons were inferred in accordance with the package pipeline. Regulon activity scores were added back to the Seurat object as a new assay. Regulons with differential activity between clusters were calculated using the “FindMarkers” function, and selected regulons for the top 20 differentially expressed regulons plotted using pheatmap^129^.

### scRNAseq and Parse workflow

Fresh whole synovium was digested into a single cell suspension, and stained for FACS as described above. For 10X, macrophages, fibroblasts, endothelial cells, and lymphocytes were sorted on the BD FACS ARIA-III. Samples were centrifuged, counted, resuspended in 43µL of serum-free media, and transported to CRUK for immediate 10X workflow. For Parse, whole live synovial cells were sorted into BSA-coated FACS tubes containing RPMI media with RNAse. Cells were then centrifuged, counted, and fixed-permeabilised as per the Parse Evercode workflow (V2.1.1), with the single modification of utilising a quarter of the recommended buffer / additive volumes to reflect the lower cell input, after discussion with Parse scientists. After the protocol, cells were counted again, and frozen in 37.5µL of cell neutralisation buffer and 2.5µL DMSO. Samples were then frozen at minus 80. The Parse Evercode WT 100k cell plate layout was designed in conjunction with Parse team advice, and samples taken on dry ice to CRUK, Cambridge. Here, the cells were thawed, counted, and loaded as per the plate design. The remaining workflow was completed by CRUK Genomics team. Quality-control and alignment was completed by Abigal Edwards (CRUK Genomics), using Parse Biosciences “split-pipe” software, aligning each sublibrary to the GRCh38 genome with ensemble annotation version 111. Cells were filtered out if they had more than 3 median absolute deviations from the median library size, median gene count per cell, and under 25% mitochondrial genes. Sample metadata was added, and it was combined with the workflow for the generation of the synovial cell atlas.

### Synovial scRNA-seq data processing and integration

In order to build the human synovial tissue atlas, publicly available datasets were assembled individually first using methods described by their respective authors^65, 66, 68–70, 93^ and the 10X and Parse sequenced scRNA-seq were also assembled as described above to be merged into a single object and then log-normalized. Highly variable genes (HVGs) were identified using the Seurat-v3 method as implemented in Scanpy (V1.10.3), selecting the top 2000 genes while accounting for batch effects across each of the individual datasets. The datasets were integrated using the scVI-module in scvi-tools (V1.2.2)^94^, the model was built with (n_layers=2, n_latent=30, gene_likelihood=“nb”),using batch keys ‘sample’, ‘dataset’ and ‘disease state’ as categorical covariates and the number of detected genes per cell as a continuous covariate. Model training was performed for 30 epochs using default optimization parameters. The integrated object was clustered by Leiden clustering at a resolution of 1.

We identified and removed Leiden cluster 3 that contained low quality fibroblast cells that were clustering together on the basis of low number of detected genes per cell due to technical variation. To annotate the broad cell types or ‘superclusters’, by making corrections to the annotations made by authors in the previous datasets manually assisted by the Cellhint tool^71^ using the ‘cellhint.harmonize’ function. After removal of the low quality cells, the dataset was then integrated using the ‘cellhint.integrate’ function by setting the batch key to the dataset parameter while considering the new assigned cell type/supercluster to adjust the neighborhood graph. Integration was conducted using the scVI latent embedding ‘X_scVI’ as the input representation, with the number of meta-neighbors set to 1 to enforce strict correspondence between matched cell populations across batches.

### Annotating the myeloid subclusters

The superclusters were then separated individually and then processed separately, to annotate the subclusters specific to the broad cell type category. The raw counts for the myeloid supercluster were log-normalized and the top 2000 HVGs were then selected using the Seurat-v3 method as implemented in Scanpy, accounting for batch effects across each of the individual datasets (‘author’). The datasets were integrated using the scVI-module in scvi-tools^94^, the model was built with (n_layers=2, n_latent=30, gene_likelihood=“nb”),using batch keys ‘sample’, ‘dataset’ and ‘disease state’ as categorical covariates and the number of detected genes per cell as a continuous covariate. Model training was performed for 100 epochs using default optimization parameters. The integrated object was clustered by Leiden clustering at a resolution of 1.

We used the (i) Alivernini myeloid dataset^65^ and (ii) the Zhang myeloid dataset^93^ as references to annotate the myeloid subclusters. For each reference, we used CellTypist^95^ (V1.6.3) to train a classifier using 2000 selected genes (feature selection enabled, 500 training iterations) and applied to the scVI-integrated dataset with majority voting enabled.

### Generating the decision matrix

The differences between the annotations in the reference datasets were subtle and in order to select the best fit annotation based on the reference as well to capture sparse cell types, we implemented an enrichment–overlap scoring framework to generate a reference–query-decision matrix. The reference here were the cell subtypes in the reference datasets and the query here refers to the Leiden clusters in the Human Synovial Tissue Atlas (HSTA).

For each reference cell type and query Leiden cluster pair, the decision score was calculated using (i) the number of overlaps between the top 50 reference celltype marker genes and the top 25 genes enriched in the query Leiden cluster (to capture the depth of gene signature being detected), (ii) the number of overlaps between the top 25 reference celltype markers and the top 10 query Leiden cluster markers (to refine the extent of genes being detected with significantly differentially expressed genes) and (iii) the median gene set enrichment score within the query cluster, computed using ‘sc.tl.score_genes’ with reference cell type-derived gene signatures of the top 100 differentially expressed genes (to fine tune the decision score).

These components were combined into a composite “decision score”:

*Decision Score* = *Median Enrichment Score* + *Overlap*(50/25) + *Overlap*(25/10)

The resulting Leiden cluster × reference cell type decision matrix for each of the reference datasets was generated, and annotation of each of the Leiden clusters was then made based on the highest decision score across the datasets.

### Annotating the osteoclast lineage cells

We found that the Leiden cluster ‘11’ in the myeloid subset showed low scores across all reference cell types in the reference datasets (**Supplementary Tables 7 and 8**). However, when mapped to the plate based scRNA seq (VASA-seq) experimental data used to identify preOCs in blood, we find that cluster 11 has the highest decision score for the preOC1 population. Therefore, cluster 11 was annotated as ‘preOCs’ (**Supplementary Table 9**). We verified this by coinciding the UMAP of the gene enrichment signal with a manually curated pre-OC specific gene list based on the plate based scRNA seq data (**Supplementary Table 10**).

To make a refined annotation, an initial manual mapping was performed by assigning Leiden clusters to candidate identities based on marker expression and differential gene expression (Wilcoxon rank-sum test). Ambiguous clusters were subsetted and reclustered at higher resolution. To further annotate osteoclast lineage populations, curated osteoclast gene signatures from osteoclast-like cells grown *in vitro* culture^41, 96^ and canonical markers (e.g., *CTSK*, *ACP5*, *MMP9*) were scored using gene set enrichment (using the top 50 DEG), and annotated as ‘OCL-1’ and ‘OCL-2’ subcluster populations.

### Trajectory analysis of the osteoclast lineage cells

The preOCs and OCLs were separated from the myeloid subset for performing trajectory analysis based on myeloid subcluster annotations. For methods requiring raw count modelling (scTour^54^), integer UMI counts were preserved. For diffusion- and PCA-based approaches (scFates^97^ and Palantir^98^), counts were library-size normalized and log-transformed.

Trajectory inference was primarily performed using scTour^54^ (V1.0.0), which uses novel deep learning architecture to perform robust inference and accurate prediction of cellular dynamics with minimal influence from batch effects. HVGs were selected using the Seurat-v3 method as implemented in Scanpy, selecting the top 1500 genes, prior to model training using scTour. The model was trained using the parameters (loss_mode=’nb’, alpha_recon_lec=0.5, alpha_recon_lode=0.5, random_state=0). Following model training, pseudotime values were extracted and stored as ptime, mixed latent space representation was stored as X_TNODE, and a vector field was inferred using learned neural ODE to model transcriptional flow. The vector field was projected onto the UMAP embedding to visualise transcriptional flow across the osteoclast differentiation trajectory.

To verify the outcome of the osteoclast lineage differentiation trajectory using scTour, it was compared against various different pseudotime calculated using different methods: (i) diffusion pseudotime^99^ which calculated using Scanpy’s function ‘sc.tl.dpt’ by setting the root cell type to ‘preOCs’ as the earliest developmental state in the trajectory, (ii) pseudotime calculated using Palantir^98^ (V1.4.10)using the default parameters by setting the root to preOCs, and (iii) pseudotime calculated using scFate^97^ (V1.1.1).

Cells were grouped into transcriptomically annotated states (preOC, OCL-1, OCL-2). Median pseudotime values per group were computed for each method, and ordering consistency across methods was evaluated.

Pairwise differences between cell states were assessed using Mann–Whitney U tests. Both two-sided and direction-aware one-sided tests were performed depending on inferred trajectory ordering. P values were adjusted for multiple testing using the Benjamini–Hochberg false discovery rate (FDR) procedure. To assess overall concordance across methods, P values for each pairwise comparison were combined using Fisher’s method, followed by FDR correction.

### Annotating the subclusters in other superclusters

The annotation of the other subclusters was conducted in a joint manner using the CellTypist model trained on the Zhang annotated subsets of their respective superclusters and then verified using DEGs of these subclusters, alongside the decision score generated using the decision matrix.

### Xenium spatial sequencing of synovial tissue samples

Appropriate synovial samples from 2 near-healthy and 2 PsA donors who had received autologous radiolabelled monocytes 24 hours before biopsy were selected. Xenium slides were equilibrated to room temperature for 30 minutes before use. Blocks were sectioned at 5µM by the histologist at Cancer Research UK, University of Cambridge, floated singly on a waterbath treated with RNase Zap (Invitrogen) before being carefully placed on the Xenium slide. Slides were dried at 42°C for 3 hours, before being stored until analysed on the Xenium Analyser (10X). The Xenium Prime 5K Human Pan-tissue and pathways panel was utilised, with cell segmentation staining. A 50 gene custom add-on specifically targeting genes relevant to synovial tissue, tissue homing, and osteoclastogenesis (**Supplementary Table 11**) was also included using the 10X panel designer and specialist advice from 10X. Decoding, segmentation and other secondary analyses were performed on the analyser as part of sample processing.

### Spatial sequencing data sample pre-processing

The output from the Xenium Analyzer (10X Genomics) for all the five samples was imported using the SpatialData package^100^ (V0.2.5).

For the sample XEN-60A, the Xenium sequencing run resulted in low number of transcript counts for particular regions of the sample as whole due to a split in the fixed block at time of sample preparation. Therefore we removed these for downstream analysis (**Supplementary Figure 7A**). K-means clustering using Scikit-learn^101^ (V1.8) on spatial coordinates using the parameters (n_clusters = 10, random_state = 52, n_init=’auto’) to identify regions to retain. The k-means clusters ‘2’ and ‘8’ coincided with transcript-counts similar to other samples and were retained for downstream analysis (**Supplementary Figure 7B**.)

### Xenium data integration and supercluster annotation

Raw count matrices were merged after filtering out low quality cells from sample XEN-60A. QC metrics were calculated on the new merged object, however to prevent cells clustering based on technical aspects, an additional parameter called ‘QC_clusters’ was considered. This was based on: the distribution between the total UMI counts and number of detected genes by applying the Gaussian Mixture Modeling (GMM; n = 4 components) on the distribution between these parameters, rather than choosing arbitrary thresholds. Cells were assigned to probabilistic QC clusters, which were stored as metadata and used as technical covariates during batch correction, to retain as many cells as possible to detect the sparse ‘preOC’ population. Cells with less than 15 detected genes and genes expressed in less than 3 cells were removed.

The dataset was normalized using ‘pp.normalize_total’ and default parameters, log-transformed, and the top 2000 HVGs were identified using sample identity as a batch key. PCA was performed, followed by an initial construction of a nearest-neighbour graph (n_neighbours=10, n_pcs=40) and compute the final UMAP embedding and Leiden clustering of the entire object.

To correct for batch effects, Harmony integration (harmonypy, v0.0.10)^102^ was applied to the PCA embeddings using both sample and QC_cluster as covariates. The Harmony-corrected embeddings were used to reconstruct the nearest-neighbour graph and compute the final UMAP embedding and Leiden clustering with default resolution of the entire object. Supercluster annotation was performed using the custom CellTypist model trained on a version of our HSTA downsampled to retain one third of every subtype population. The annotation was verified by cross-referencing the DEG of each of the superclusters of the Xenium Dataset against the PanglaoDB database^103^. Each of the superclusters were then separated and the subcluster annotation was performed for the myeloid populations.

### Myeloid subcluster annotation

Raw counts for the myeloid population were renormalized to 10000 counts per cell, log-transformed, and HVGs (800 genes) were identified using sample identity as a batch key. Neighbourhood graphs and UMAP embeddings were generated using 10 principal components. Leiden clustering (resolution=0.5) was performed, and cluster-specific marker genes were identified using the tl.rank_gene_groups function. Clusters expressing non-myeloid markers were removed (Leiden clusters:’5’, ‘9’, ‘10’ and’11’) (**Supplementary Figure 7D**), and the raw counts matrix for these cells were then generated.

In order to correct for batch effects, Harmony integration was applied to the PCA embeddings using both ‘sample’ and ‘QC_cluster’ as covariates. The Harmony-corrected embeddings were used to reconstruct the nearest-neighbour graph and compute the final UMAP embedding and Leiden clustering at a resolution of 0.8 for the myeloid population.

The Leiden clusters were then annotated using the decision score matrix matched against the myeloid cell types in the HSTA (**Supplementary Figure 7E**). Leiden clusters ‘6’ and ‘17’ showed no enrichment for the majority of cell types, however they displayed high levels of expression of genes specific to preOC cells characterised in the HSTA and that of the *DCSTAMP* gene which was used to identify preOC in blood (**Supplementary Figure 7F, left panel)**. As the majority of the top 100 preOC-specific genes from the HSTA were highly expressed in these clusters relative to other myeloid clusters, we annotated these clusters ‘preOC’ in the Xenium spatial dataset. 53 genes are visualised for representative purposes (**Supplementary Figure 7F, right panel**).

Xenium Explorer^104^ (V4.0) was used for the spatial visualization of clusters and superclusters.

### Radiolabelling and imaging

#### *In vitro* experiments

160ml of blood was taken from healthy volunteers using a 19G needle and into EDTA collection tubes (BD, 367525). Cells were processed into PBMCs as described above. Monocytes were isolated as described above. 550MBq of Tc^99m^ and sterile hexamethylpropyleneamine oxime (HMPAO) was obtained from Cambridge University Hospitals NHS Foundation Trust radiopharmacy. Radioactive materials were transported to the Department of Pathology by the University of Cambridge Radiation Protection Services in appropriate lead vessels in accordance with UK law. Tc^99m^ was drawn up with a syringe and mixed with HMPAO for reconstitution. Tc^99m–^HMPAO was then immediately added to the cell pellet and incubated for 15 minutes. Cells were then washed in FACS buffer, and the pelleted cells resuspended in Cryostor CS10 and immediately frozen in lead vessels. The Tc^99m–^labelled cells were then allowed to completely decay, before being thawed, and cultured into moMacs as described above. Cells were removed from plates using accutase, and stained for mass cytometry as described above.

#### Cell isolation and radiolabelling

Cells were isolated from peripheral venous blood using discontinuous plasma-Percoll gradients and radiolabelled, as previously described^105, 106^. Monocytes were purified using clinical-grade CD14 microbeads (CliniMACS; Miltenyi Biotec 170-076-705). Purified monocytes were radiolabelled with Tc^99m–^hexamethylpropylene amine oxime (HMPAO) for imaging studies.

#### In vivo imaging

Radiolabelled monocytes were reinfused as an intravenous bolus into healthy or psoriatic volunteers. For imaging, participants were positioned in a dual-headed SPECT/CT camera (GE Discovery 670, GE Healthcare), fitted with low-energy high-resolution, parallel-hole collimators. From the point of bolus intravenous injection of autologous Tc^99m–^labelled monocytes, the counts in the chest and upper abdomen were recorded as previously described over 40 minutes. Following this, sequential planar imaging of the knees, feet and ankles, and hands and wrists were performed. Areas were visualised for 20 minutes each, with imaging performed at 2-, 4-, and 24-hours. A radioactive marker was used to mark the right-hand side of images.

Images were generated as described above, and exported to Cambridge University Hospitals NHS Foundation Trust. Images were loaded into Xeleris (GE Healthcare) by senior scientists in the nuclear medicine team. Regions of interest were drawn over relevant areas on all scans. Monocyte uptake to the region was quantified by a nuclear medicine physicist. Lung, liver, and spleen uptake was quantified. Data was provided as .csv files.

### Statistical analysis

All relevant statistical analyses were performed in GraphPad Prism v9.2.0 (GraphPad) or R. They are indicated in the relevant figure legends throughout. Significance was reported when corrected p-values were under 0.05. For flow and mass cytometry data, quantification and geometric median fluorescence intensity (gMFI) data were exported from FlowJo and analysed in GraphPad Prism.

## Data availability

All the RNA sequencing data generated in this study are being uploaded to EMBL-EBI ArrayExpress. Codes used for all the analyses will also be shared.

Data are available under the terms of the Creative Commons Attribution 4.0 International license (CC-BY 4.0).

## Acknowledgements

We thank the Flow Cytometry Core Facility at the Department of Pathology, Richard Grenfell (CRUK, flow core, Cambridge) and the CRUK Genomics Core, Cambridge for their assistance. We thank Dr Sarah McDonald (Cambridge University Hospitals NHS Foundation Trust) for her kind histopathological assessment of synovial tissue samples. We thank all donors who participated in this study and hospital staff.

We thank Mariola S Kurowska-Stolarska and Iain McInnes (University of Glasgow) for scientific discussion and guidance.

We thank Narendra Meena for scientific discussion and for assistance with SCENIC transcription factor visualisation.

## Author contributions

Conceptualization, JPH, CS, N.McG; methodology, JPH, AS, AL, QL, NF, DG, FL, BA, DJ, NS, AL, CS, N.McG; formal analysis, JPH, AS, QL, NM, FL, NF, AL, N. McG; investigation, JPH, AS, QL, NM, FL, NF, AL, N. McG; writing – original draft, JPH, AS, N.McG; visualization, JPH, AS, AL, QL, N.McG; supervision, N.McG, AL and CS.; funding acquisition, N.McG, CS, JPH.

